# Convergent loss of the necroptosis pathway in disparate mammalian lineages shapes virus countermeasures

**DOI:** 10.1101/2021.06.08.447400

**Authors:** Ana Águeda-Pinto, Luís Q. Alves, Fabiana Neves, Grant McFadden, Bertram L Jacobs, L. Filipe C. Castro, Masmudur M. Rahman, Pedro J. Esteves

**Affiliations:** CIBIO/InBio-Centro de Investigação em Biodiversidade e Recursos Genéticos, Universidade do Porto, Vairão, Portugal; Departamento de Biologia, Faculdade de Ciências, Universidade do Porto, Porto, Portugal; CIIMAR/CIMAR, Centro Interdisciplinar de Investigação Marinha e Ambiental, Universidade do Porto, Matosinhos, Portugal; Center for Immunotherapy, Vaccines and Virotherapy (CIVV), The Biodesign Institute, Arizona State University, Tempe, USA; School of Life Sciences Center for Immunotherapy, Vaccines and Virotherapy, Biodesign Institute, Arizona State University, Tempe, USA; CITS-Centro de Investigação em Tecnologias da Saúde, IPSN, CESPU, Gandra, Portugal

## Abstract

Programmed cell death is a vital process in the life cycle of an organism. Necroptosis, an evolutionary restricted form of programmed necrosis, contributes to the innate immune response by killing pathogen-infected cells. This virus-host interaction pathway is organized around two key components: the receptor-interacting protein kinase 3 (RIPK3), which recruits and phosphorylates the mixed lineage kinase-like protein (MLKL), thus inducing cellular plasma membrane rupture and cell death. Critically, the presence of necroptotic inhibitors in viral genomes validates necroptosis as an important host defense mechanism. Here, we show, counterintuitively, that in different mammalian lineages of mammalian, central components of the necroptotic pathway, such as *RIPK3* and *MLKL* genes, are deleted or display inactivating mutations. Frameshifts or premature stop codons are observed in all the studied species of cetaceans and leporids. In carnivores’ genomes, the *MLKL* gene is deleted, while in a small number of species from afrotheria and rodentia premature stop codons are observed in *RIPK3* and/or *MLKL*. Interestingly, we also found a strong correlation between the disruption of necroptosis in leporids and cetaceans and the absence of the C-terminal domain of E3-like homologs (responsible for necroptosis inhibition) in their naturally infecting poxviruses. Overall, our study provides the first comprehensive picture of the molecular evolution of necroptosis in mammals. The loss of necroptosis multiple times during mammalian evolution highlights the importance of gene/pathway loss for species adaptation and suggests that necroptosis is not required for normal mammalian development. Moreover, this study highlights a co-evolutionary relationship between poxviruses and their hosts, emphasizing the role of host adaptation in shaping virus evolution.

## Introduction

Sensing of viral pathogens by the host cells is critical for animal survival. Thus, a variety of molecular responses, including the induction of inflammatory cytokines, chemokines and interferons, as well as the activation of cell-death pathways that provide clearance of pathogen-infected cells. Although apoptosis has long been considered a critical clearance mechanism to control viral spread, caspase-independent cell death, or programmed necrosis, has recently emerged as an alternative death pathway that dominates under specific conditions (Xia et al. 2020).

Necroptosis is an inflammatory form of regulated necrosis that acts as an alternative host defense pathway during some viral infections and plays a major role in the killing and removal of pathogen-infected cells (Upton and Chan 2014; Nogusa et al. 2016; Xia et al. 2020). Activation of necroptosis follows an intracellular signaling cascade that is dependent on the receptor-interacting serine/threonine-protein kinase 3 (RIPK3) and its substrate, the mixed lineage kinase like protein (MLKL) downstream of death receptors (DRs) and pattern-recognition receptors (PPRs) (Fig. 1) (Sun et al. 2012; Moerke et al. 2019). Several pathway-specific adaptor proteins that contain a RIP homotypic interaction motif (RHIM-domain) can activate RIPK3-induced necroptosis (Fig. 1). For example, when there is an interference or loss of function of caspase-8, the induction of necroptosis through the use of DRs results in the recruitment of RIPK1, which subsequently exposes its RHIM-domain to recruit RIPK3 (He et al. 2009; Zhang et al. 2009; Upton et al. 2010). Apart from RIPK1, the TIR-domain-containing adaptor-inducing IFN β (TRIF), an essential protein downstream of Toll-like receptor (TLR)3/4 and the Z-DNA binding protein (ZBP1), also directly activate RIPK3 (He et al. 2011; Upton et al. 2012) (Fig. 1). Exposure of RIPK3 to a RHIM adaptor (RIPK1, TRIF or ZBP1) is a crucial step in the initiation of necroptosis, as these proteins activate the downstream executor of necroptosis MLKL that destabilizes the plasma membrane integrity leading to cell swelling followed by membrane rupture of infected cells and release of damage-associated molecular patterns (DAMPs) (Kaiser et al. 2013; Upton and Chan 2014). Thus, necroptosis provides a critical extra defense mechanism against pathogen infection, facilitating the elimination of virus-infected cells before the production of progeny virions. The importance of necroptosis for host defense is further supported by the identification of viral inhibitors of necroptosis, like is the case of Vaccinia virus (VACV) E3 protein and the murine cytomegalovirus (MCMV) M45 protein (Upton et al. 2012; Kaiser et al. 2013).

**Fig. 1.**
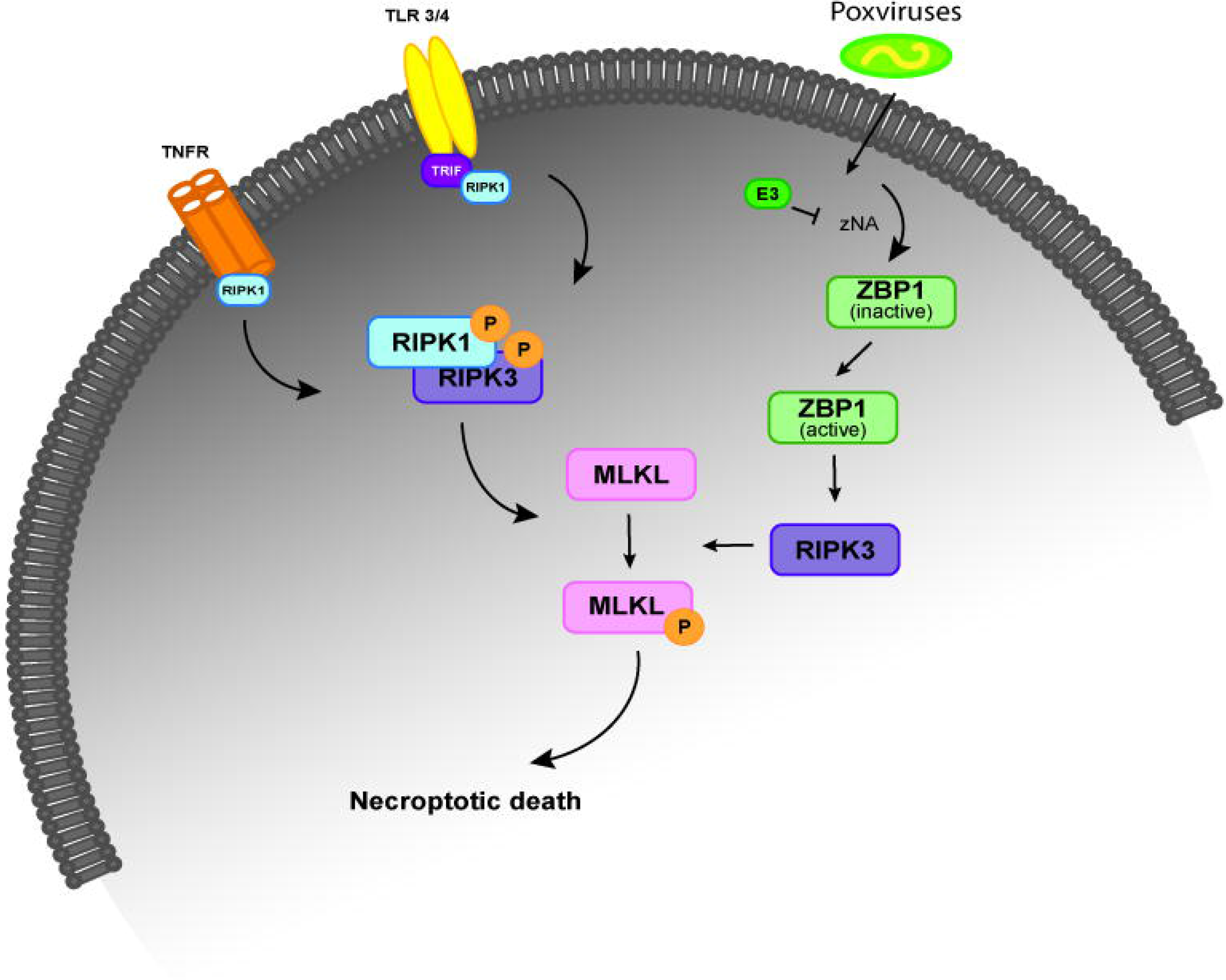
The necroptosis signaling pathway. Simplified schematic representation of the necroptosis signaling pathway upon stimulation of the TNFR, TRF3/4 and infection by poxviruses. All of these necroptosis-inducing signals converge on the kinase RIPK3, which is activated through the homotypic interaction with RIPK1 or other RHIM-containing proteins, such as TRIF and DAI. When the activity of caspase-8 is inhibited, binding of TNF to TNFR1 leads to the phosphorylation and activation of RIPK1 that binds to RIPK3 through their RHIM domains to form a protein complex (necrosome). Activated RIPK3 recruits MLKL that oligomerizes and translocates to the plasma membrane to cause necroptosis. In TLR3- and TLR4-induced necroptosis, TRIF is required for the activation of RIPK3. ZBP1 is required for the activation of RIPK3 in response to the presence of Z-form nucleic acids. In VACV-infected cells, the poxviral E3 protein binds to VACV-induced Z-form nucleic acid, preventing RIPK3-induced necroptosis. Abbreviations: TNFR, tumor necrosis factor receptor; TLR 3/4, toll-like receptor; TRIF, TIR-domain-containing adaptor-inducing IFN β; RIP, receptor-interacting protein kinase; ZBP1, Z-DNA binding protein; MLKL, mixed-lineage kinase domain like.

Necroptosis has a major role in protecting cells against viral infection (Upton and Chan 2014; Nogusa et al. 2016; Xia et al. 2020). However, despite the recent advances to understand the molecular regulation of this unique pathway, it is still unclear whether the necroptotic cell death pathway acts as a universal cell death program in mammals. Previous studies suggested that components of necroptosis are absent in the genomes of extant birds and marsupials. Interestingly, within the Mammalia class, it was previously reported that order Carnivora lost the *MLKL* gene (Dondelinger et al. 2016). Taken together these reports suggest some degree of plasticity in the conservation of necroptosis responses. Here, we address the molecular evolution of the necroptotic pathway in multiple mammalian lineages. We show that during mammalian evolution, necroptosis was convergently inactivated several times in mammalian lineages. Remarkably, we also report that mammalian orders that lost the necroptotic pathway display infection episodes by poxviruses that have lost the ability to inhibit this pathway, showing a co-evolutionary relationship between host adaptation in shaping virus evolution.

## Results

Necroptosis has a significant role in protecting cells against viral infection (Upton and Chan 2014; Nogusa et al. 2016; Xia et al. 2020). However, despite recent advances to understand the molecular regulation of this unique pathway, it is still unclear whether the necroptotic cell death pathway acts as a universal cell death program in mammals. Here, taking advantage of genomic collection databases and the use of Leporid samples (see Methods section for more information), the goal is to better understand the molecular evolution of the necroptotic pathway in different mammalian lineages.

### RIPK1 protein is under evolutionary conservation in mammals

Human RIPK1 is a multidomain protein composed of an N-terminal Ser/Thr kinase, a C-terminal death binding domain that mediates binding to DRs and an intermediate domain that includes a K377 ubiquitination site and an RHIM motif that binds to other RHIM-containing proteins (He and Wang 2018). Due to the importance of RIPK1 as an adaptor molecule, previous phylogenetic analysis suggested that RIPK1 probably emerged in the ancestor of vertebrates (Dondelinger et al. 2016). In accordance, our search detected the presence of *RIPK1* homologues in all the studied mammalian lineages (Fig. 2A and S Appendix 1, 2 and 3).

**Fig. 2.**
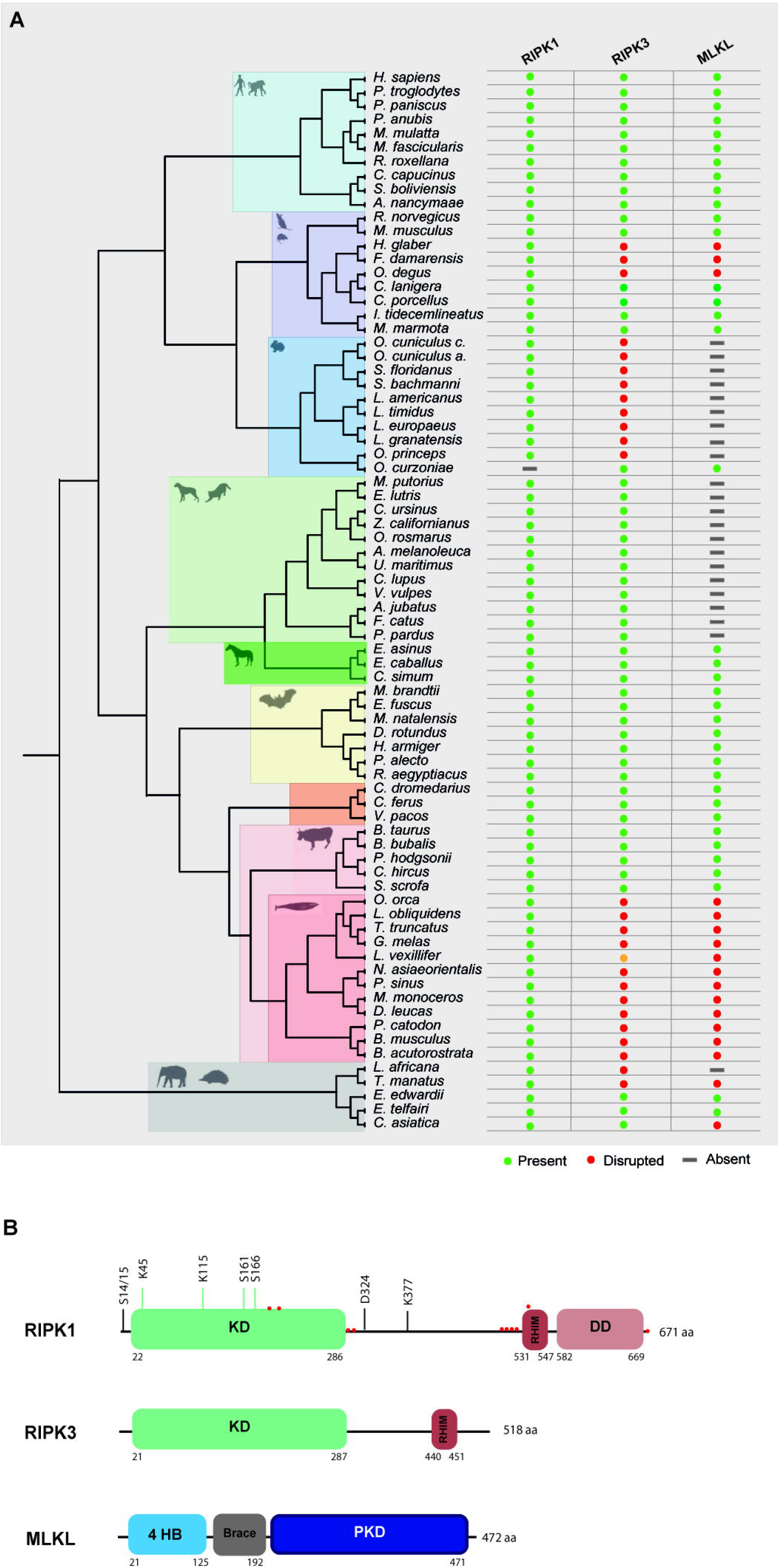
Evolution of RIPK1, RIPK3 and MLKL in different mammalian lineages. **A)** Phylogenetic tree showing the independent lineages that lost necroptotic core proteins (RIPK3 and MLKL) during evolution. Green circles represent genes that are present in the studied species, red circles represent genes that are disrupted, yellow circles represent genes that have incomplete assemblies and grey rectangles indicate that genes were not found in those species genomes. **B)** A schematic diagram of RIPK1, RIPK3 and MLKL domains. RIPK1 contains an N-terminal kinase domain (KD), an intermediate domain with a RIP homotypic interaction motif (RHIM), and a C-terminal death domain (DD). The phosphorylation and ubiquitination sites are indicated above the RIPK1 domains. Red circles represent residues that are under positive selection. RIPK3 contains a KD and a RHIM domain. MLKL is composed of an N-terminal bundle four-helix bundle (4HD) domain that is regulated by the C-terminal pseudokinase domain (PKD).

To look for evidence of potential selection pressures acting on the different domains of RIPK1 protein, we used the dataset of mammalian sequences mentioned above and implemented an ML approach, by using the Datamonkey software (see Methods section for more information). For most protein-coding genes, the rate between nonsynonymous and synonymous substitutions (dN/dS) is a measure of natural selection, with positive selection (dN/dS > 1) acting against the common genotype (Kosiol et al. 2008). In this study, we deduced ten sites that reflect strong positive selection pressure in RIPK1, while more than 200 amino acids were under negative selection (Fig. 2B and S Appendix 4). As seen in Fig. 2B, residues identified as being under positive selection fall within or very close to the kinase domain (4 residues), the RHIM domain (5 residues) and the death domain (1 residue) of RIPK1 (residues under positive selection are marked as red circles). The N-terminal kinase domain is known to present several essential residues for phosphorylation and ubiquitination (Ser14/15, 20, 161 and 166 and Lys115 and 163), regulation of necroptosis and RIPK1-dependent apoptosis (Mifflin et al. 2020). Interestingly, the codons under positive selection fall only at the end of the N-terminal domain. A considerable portion of the negatively selected sites fall in the critical regions for phosphorylation and ubiquitination (S Appendix 4), suggesting that the beginning of the RIPK1 protein is under strong purifying selection. The same was also observed for the rapidly evolving sites of the RHIM domain, which were grouped only at the beginning of this domain (Fig. 2B). It was previously shown that the RHIM domain has a crucial conserved core motif of 12 amino acids that resides at the end of this domain (Sun et al. 2002; Li et al. 2012). Indeed, changing four consecutive amino acids to alanine within this core region abrogates interaction between RHIM domains and, as a consequence, necroptosis (Sun et al. 2002; Li et al. 2012). Our findings that the positively selected residues did not overlap with the core motif of 12 amino acids further support the importance of the conservation of this region. Interestingly, in the RIPK1 death domain that is known to mediate homodimerization as well as heterodimerization with other DD-containing proteins, such as FADD, TNFR1 and Fas (Meng et al. 2018; Mifflin et al. 2020), no positively selected sites were found, with most of the domain being under negative pressure (S Appendix 4). Collectively, our results show signatures of positive selection occurring at the end of the N-terminal kinase domain and at the beginning of the RHIM domain of RIPK1 proteins. Interestingly, these residues do not overlap with domains known to be fundamental for RIPK1-dependent apoptosis and necroptosis, suggesting that these domains might be under evolutionary conservation and possibly functional constraint for the studied mammals.

### Convergent erosion of RIPK3 and MLKL in mammalian lineages

RIPK3 and MLKL form the core of the necroptotic machinery and both are, as a consequence, important for necroptosis induction in mice and humans downstream of PRRs and DRs (Sun et al. 2012; Moerke et al. 2019; Xia et al. 2020). To further understand the evolutionary history of the necroptotic pathway in mammals, we performed detailed sequence and phylogenetic analyses for RIPK3 and MLKL homologous proteins (S Appendix 3). Our screens for *RIPK3* and *MLKL* genes revealed evidence of pseudogenization in five mammalian lineages: order rodentia, lagomorpha, carnivora, cetacean and in the superorder afrotheria (Fig. 2A). Given the fact that *MLKL* pseudogenes have been previously identified in carnivores, they will not be discussed in detail here (Dondelinger et al. 2016).

### Variation of the Necroptotic pathway within Afrotherian and Rodent families

In afrotherian and rodent lineages, we found that the necroptotic pathway was missing in some species (Fig. 2A). In rodents, two species from the Bathyergidae family and one species from the Octodontidae family presented early stop codons in both RIPK3 and MLKL, resulting in the disruption of necroptosis. In the naked mole-rat (*Heterocephalus glaber*), RIPK3 presented a premature stop codon in exon 6 resulting in a shorter version of this protein (S Appendix 5). In the common degu (*Octodon degus*) and in the damaraland mole-rat (*Fukomys damarensis*), both RIPK3 and MLKL proteins presented several premature stop codons (S Appendix 5). However, disruption of these proteins appears to have occurred in an independent way, rather than in a common ancestral. Given the fact that *RIPK3* and *MLKL* present signs of pseudogenization, it is expected that in the naked mole-rat, common degu and damaraland mole-rat necroptosis is disrupted. Our studies also revealed that species from Afrotherian families, including the African bush elephant (*Loxodonta africana*, family Elephantidae) and the Cape golden mole (Chrysochloris asiatica, family Chrysochloridae) presented early stop codons in RIPK3 and MLKL, respectively, while the the West Indian manatee (*Trichechus manatus*, family Trichechidae) present early stop codons in both genes (S Appendix 5). However, our results also show that the Cape elephant (*Elephantulus edwardii*, family Macroscelididae) and the lesser hedgehog tenrec (*Echinops telfairi*, family Tenrecidae) present intact copies of the *RIPK3* and *MLKL* genes, indicating that RIPK3 and MLKL were present in early stages of Afrotheria evolution, but must have been lost later in specific lineages, resulting in the existence of alternative modes of necroptosis inactivation.

### The Necroptotic pathway is disrupted in Lagomorphs

The Order Lagomorpha is divided into two families, Ochotonidae and Leporidae, which diverged around 30–35 Mya (Melo-Ferreira et al. 2015). While Ochotonidae is only composed of one extant genus, *Ochotona*, the Leporidae family includes 11 genera, including *Lepus*, *Sylvilagus* and *Oryctolagus* (Ge et al. 2013). Using the methods described previously, we were only able to identify *RIPK3* and *MLKL* transcripts for plateau Pika (*O. curzoniae*), while no *RIPK3* and *MLKL* transcripts were found for the European rabbit (*O. cuniculus*). For the American Pika (*O. princeps*), incomplete genome assemblies in the vicinity of the *RIPK3* and *MLKL* regions made retrieving the sequence of these genes impossible. By evaluating *RIPK3* gene from human and mouse and its genomic context, we were able derive a partial *RIPK3* nucleotide sequence from the European rabbit genome, which displays a frameshift mutation caused by the insertion of a single nucleotide. It is well known that accurately detecting gene-inactivation mutations in these alignments poses a number of challenges like, for example, sequencing errors and cases of assembly incompleteness. For this reason, we assessed the accuracy of our database prediction by sequencing that same genomic region in different Leporid species, representative of different genera (*Lepus*, *Sylvilagus* and *Oryctolagus*). From the obtained results, we confirmed the insertion of 1 nucleotide (+G, exon 3) not only in the European rabbit RIPK3, but also in species from genus *Lepus* and *Sylvilagus*, suggesting that disruption of *RIPK3* gene occurred in a common ancestral and was maintained throughout Leporid evolution (Fig. 3A). During necroptosis, activated RIPK3 phosphorylates and activates MLKL, which results in its recruitment and oligomerization in the plasma membrane leading to rupture and cell death (Sun et al. 2012; Xia et al. 2020). Interestingly, and despite all of our efforts, no *MLKL* transcripts or MLKL protein accumulation were found in any of the studied Lagomorphs (data not shown). Detailed analysis of the upstream and downstream MLKL flanking genes in both human and mouse genomes reveal that MLKL resides between the ring finger and WD repeat domain 3 protein (*RFWD3*) and fatty acid 2-hydroxylase (*FA2H*) genes (Fig. 4). Accordingly, although there are no gaps or incomplete genomic assemblies surrounding that region in the European rabbit genome, we were not able to retrieve a complete or partial *MLKL* gene, suggesting once more that this gene is not present in these mammals. Together, our results suggest that the studied leporid species are deficient in the core proteins of the necroptotic pathway, and that RIPK3 inactivation occurred at the stem Leporid branch and was maintained during evolution.

**Fig. 3.**
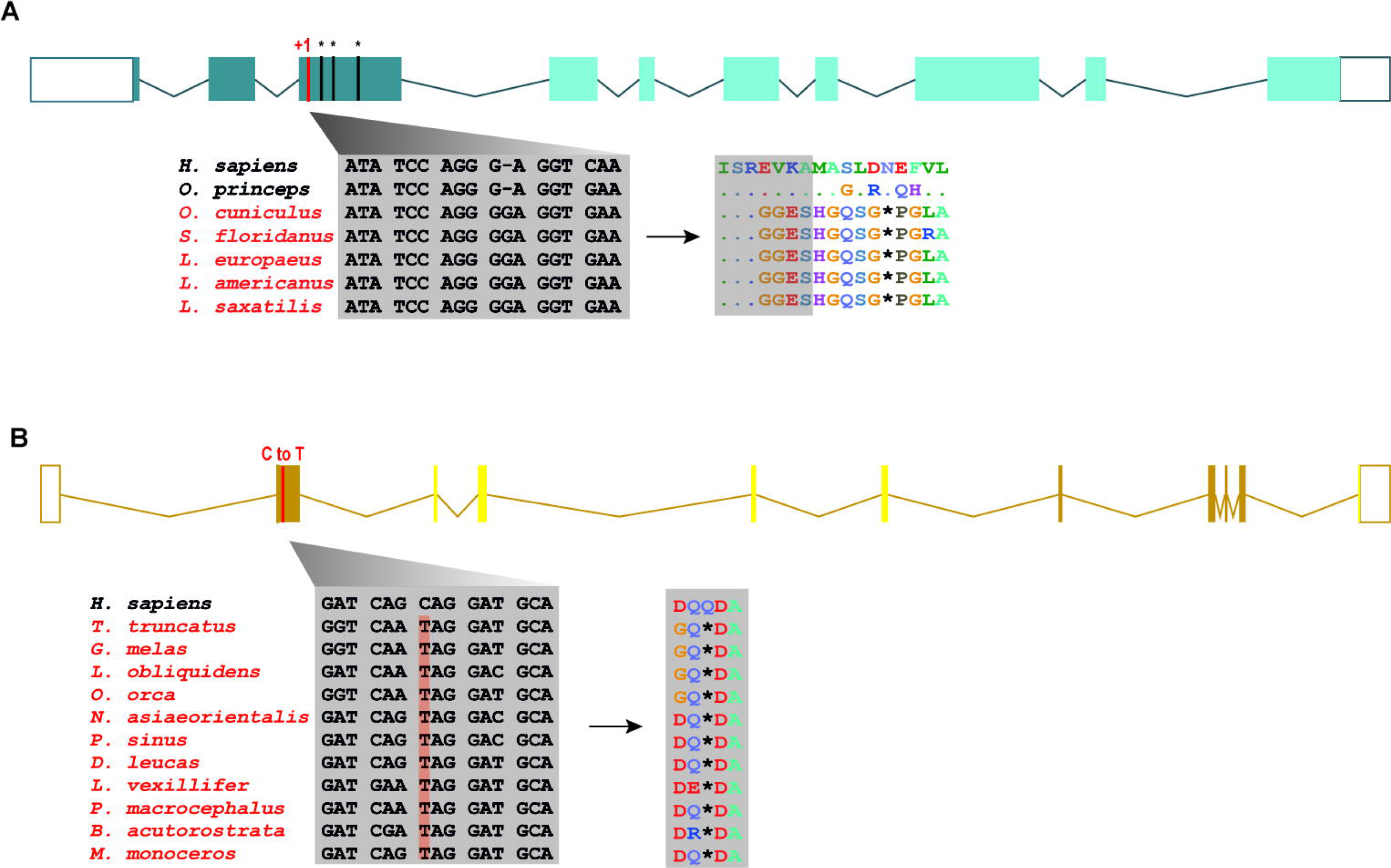
Loss of *RIPK3* and *MLKL* genes in the steam lineage of Leporids and Cetaceans. **A)** Genomic analysis of the first tree exons from Leporids (marked as dark green). In Leporids, RIPK3 was lost as a result of a shared insertion (+G) in the third exon that resulted in the appearance of several premature stop codons. **B)** A point mutation (C to T) in all the studied cetacean species indicates that MLKL inactivation occurred in Cetacea steam lineage. Moreover, 9 out of the 11 studied species (excluding *B. acutorostrata* and *M. Monoceros*) lost exon 2, 3, 4 and 5 throughout evolution (represented by faint yellow). Premature stop codons are represented by an asterisk (*).

**Fig. 4.**
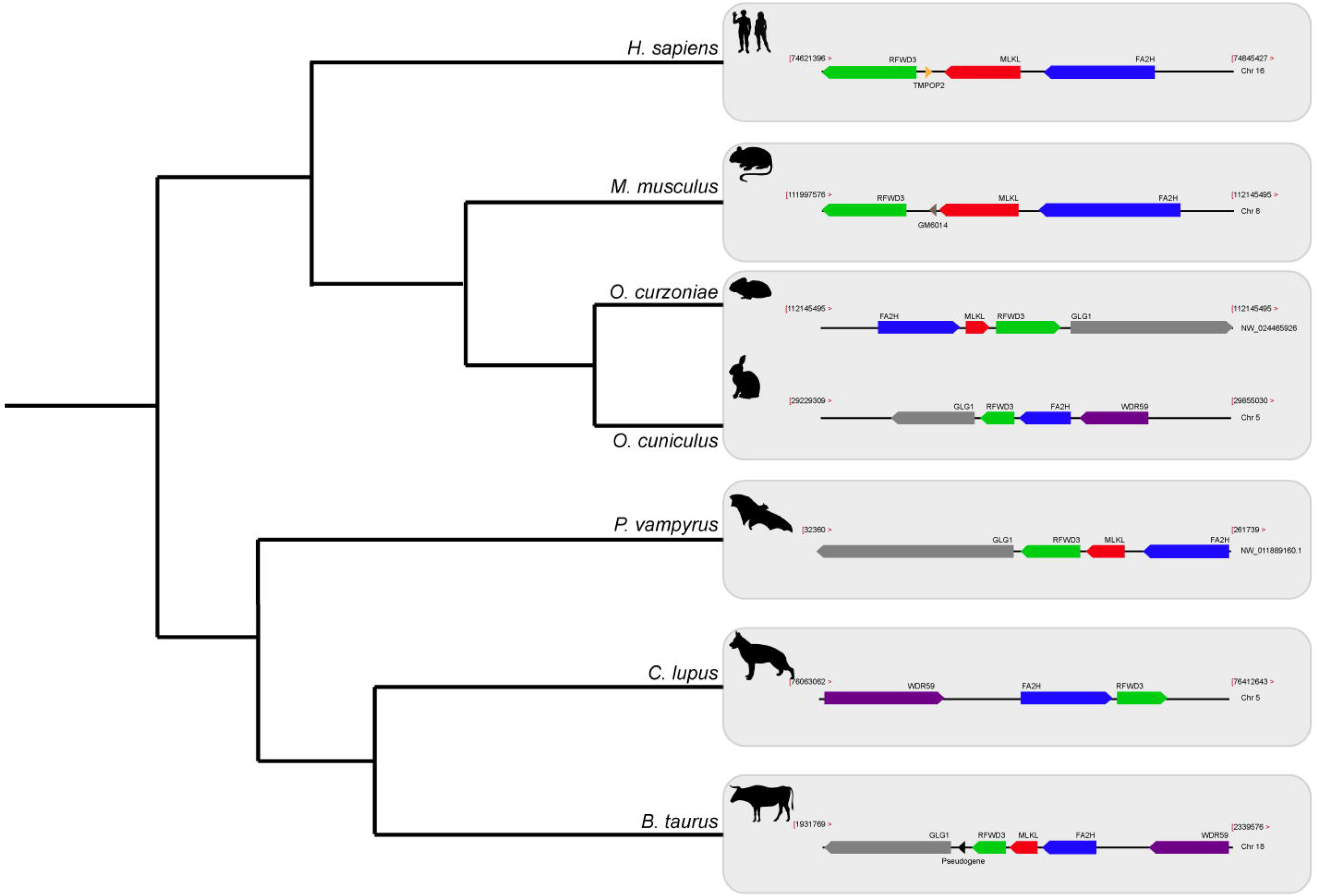
Gene synteny of the genome regions containing MLKL gene in different mammals. Genomic regions containing the *MLKL* gene or its flanking genes in *H. sapiens*, *musculus*, *O. curzoniae*, *O. cuniculus*, *P. vampyrus*, *C. lupus* and *B. taurus*. Horizontal lines indicate chromosome fragments and coloured arrows identify genes and their orientation in the genome. Orthologous genes are indicated in the same colour and their names are indicated above/below. Black arrows indicate the presence of pseudogenes. Abbreviations: RFWD3, ring finger and WD repeat domain 3; MLKL, mixed-lineage kinase domain like; FA2H, fatty acid 2-hydroxylase; GLG1, golgi glycoprotein 1; WDR59, WD repeat domain 59; *TMPOP2*, thymopoietin pseudogene; *GM6014*, ubiquitin-40S ribosomal protein S27a pseudogene; *LOC788457*, translationally-controlled 1 pseudogene.

### Inactivation of necroptosis components in Cetacea

Modern Cetacea comprises Mysticete (or baleen whales) and Odontocete (or toothed whales) and are the most specialized and diversified group of mammals (McGowen et al. 2020). Comparative analysis of cetacean genomes has already provided important insights into the unique cetacean traits and aquatic specializations (Sharma et al. 2018; Huelsmann et al. 2019; Kawasaki et al. 2020). For our screen, Odontocetes were represented by 12 species belonging to five different families (Delphinidae, Phocoenidae, Monodontidae, Lipotidae and Physeteridae), and Mysticetes were represented by the common minke whale from Balaenopteridae family. In Cetacean species, the disruption of *RIPK3* occurred at different positions depending on the studied species: a frameshift mutation was identified in exon 6 of two Delphinidae species, as well as in exon 8 of two Phocoenidae species, one Monodontidae species and one Balaenopteridae species. There was also evidence of two species (one species from Monodontidae and another from Lipotidae families) with RIPK3 pseudogenes based on the presence of a stop codon located in exon 2 (S Appendix 6). Interestingly, while Cetacean RIPK3 inactivation appears to be a result of different mutations depending on the studied species, our results show a shared mutation in the exon 1 of *MLKL* in all cetacean species (Fig. 3B). Moreover, this premature stop codon leads to the absence of exon 2, 3, 4 and 5 in most cetacean species, which very likely results in this gene inactivation. Given the presence of an inactivating mutation that is shared between mysticetes and odontocetes, the most parsimonious hypothesis suggests that they occurred before the split of these two clades in the common ancestral branch of Cetacea.

### Diversity among the poxvirus encoded E3-like necroptosis antagonists

Previously, it was shown that the N-terminus of VACV E3 competes with ZBP1 for binding to virus-induced Z-nucleic acid, being a key component to inhibit the action of IFN and induction of necroptosis (Koehler et al. 2017) (Fig. 1). E3-like encoded proteins are composed of a carboxy (C)-terminal double-stranded RNA binding domain (dsRNA-BD) and an amino (N)-terminal Z-nucleic acid-binding domain (zNA-BD) (Kim et al. 2003; Kim et al. 2004) (Fig. 5A). Given the importance of the N-terminus region from VACV E3 protein against virus infection, we hypothesize that poxviruses that lack this region in their E3 homologs can still successfully replicate in their natural host because of a compromised necroptotic pathway. Among the E3L proteins, E3 from VACV is the best studied protein. However, E3 homologs can be found in orthopoxviruses, clade II poxviruses and parapoxviruses (Kim et al. 2003; Kim et al. 2004). Recently, the genome characterization of CePV-TA identified two novel E3L homologs: CePV-TA-20 and CePV-TA-21 (Rodrigues et al. 2020).

**Fig. 5.**
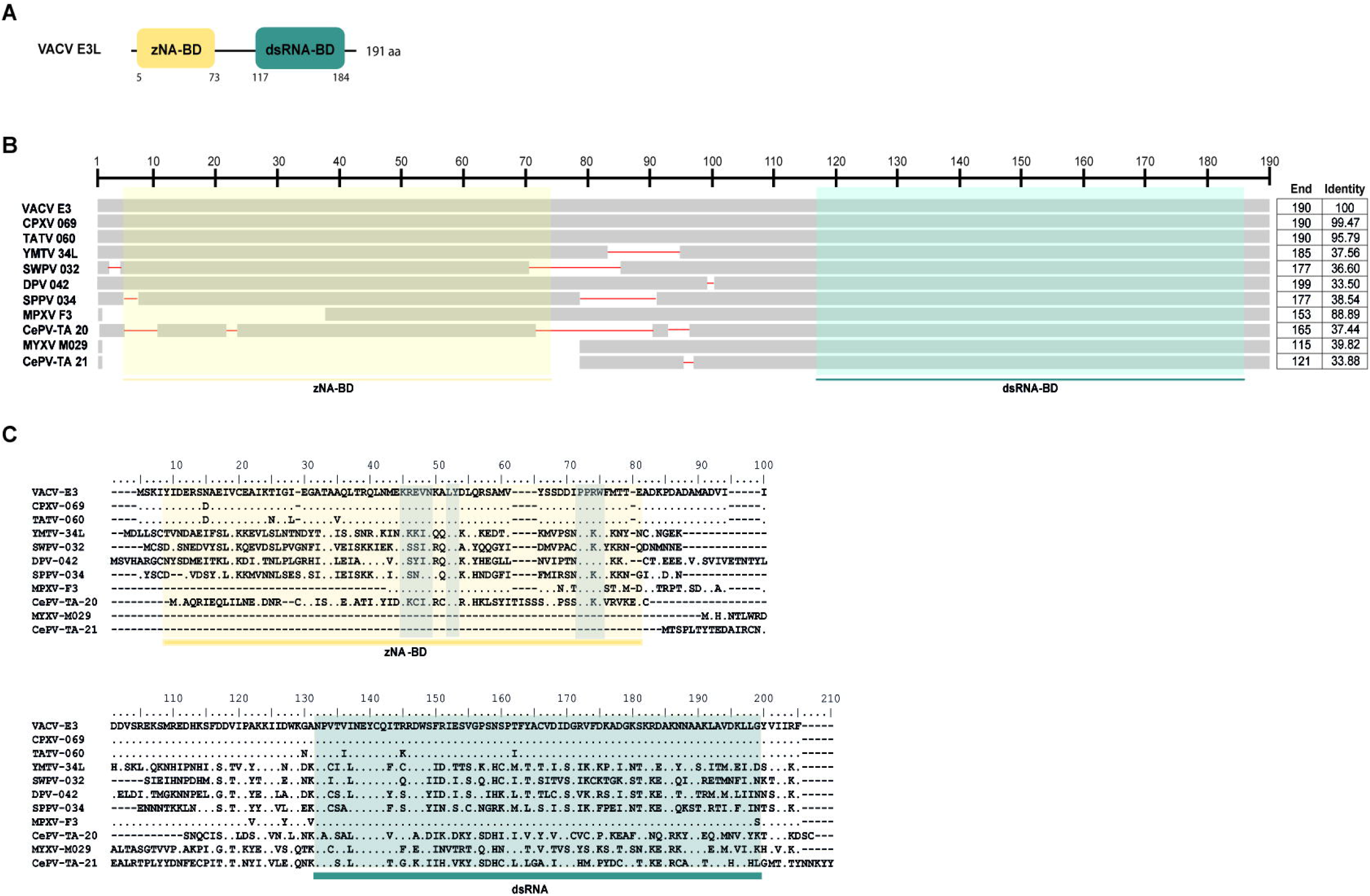
Protein sequence alignment of E3L proteins. **A)** Schematic diagram of VACV E3 protein binding domains: yellow box represents the zNA-BD and blue box represents the dsRNA-BD. The same color scheme is used in B and C. **B)** E3L homologues from 11 poxviruses (VACV E3, Cowpoxvirus (CPXV) 069, Tateropoxvirus (TATV) 060, Yaba monkey tumor virus (YMTV) 034, swinepoxvirus (SWPV) 34L, Deerpoxvirus (DPV) 042, Sheeppoxvirus (SPPV) 034, Monkeypoxvirus (MPXV) F3, Myxoma virus (MYXV) M029 and Cetaceanpoxvirus (CePV) CePV-TA-20 and 21) were used to perform a schematic alignment using COBALT program from the NCBI platform. Length of each E3L homologue as well as their identity to VACV E3 proteins are shown in the column to the right. **C)** Amino acid sequence comparison of 11 different members of the E3L family including VACV E3, TATV 060, YMTV 034, SWPV 34L, DPV 042, SSPPV 034, MPXV F3 and MYXV M029 and CePV-TA 20 and 21. Conserved areas known to bind to zNA are shown in grey boxes.

Our analysis on 11 different E3 homologues revealed that these are highly divergent: while CPXV 069 (Cowpoxvirus E3 homolog) and TATV 060 (Tateropoxvirus E3 homolog) presented identities of >90% to VACV E3, E3L homologs from poxviruses like the Deerpoxvirus, Sheeppoxvirus and Yaba monkey poxvirus presented less than 40% identity (Fig. 5B). Analysis of the two newly identified E3 homologs from CePV-TA shows that both present low identity to VACV E3, with CePV-TA-20 and CePV-TA-21 proteins only presenting sequence identity of 37 % and 34 %, respectively (Fig. 5B). It is known that at the amino acid level, the C-terminal of E3-like proteins display a higher level of sequence similarity than the N-terminal domain (Rahman and McFadden 2020). Accordingly, the dsRNA-BD domain from CePV-TA-20 and CePV-TA-21 proteins also display a higher level of sequence similarity compared to other E3 homologs (Fig. 5C), suggesting that in CeTV this domain might also target conserved antiviral dsRNA-activated pathways. Similar to what is observed for MPXV and MYXV E3 homologues, both E3 homologs from CePV-TA present incomplete or disrupted N-terminal zNA-BDs. As shown in Fig. 5C, CePV-TA-20 is missing 20 amino acids in its N-terminal domain. However, this region still retains the conserved LY and PPXW motifs, as well the basic KKCINR motif (Fig. 5C), residues known to contact with Z-NA (Kim et al. 2003). Interestingly, MPXV F3 protein, also lacking 37 amino acids from the N-terminal domain, is not able to compete with ZBP1 and inhibit sensing (unpublished data), even though it retains the key residues important for binding to Z-NA (Fig. 5C). While CePV-TA-20 and F3 proteins contain an incomplete zNA-BD, M029 and CePV-TA-21 proteins are missing most of their N-terminal zNA-BD (Fig. 5B and C), suggesting a total inactivation of this domain and a loss of function regarding the inhibition of ZBP1-dependent necroptosis. Overall, our results show that the novel CePV-TA presents two E3L homologues that, like E3L homologues from MPXV and MYXV, present incomplete or disrupted N-terminal zNA-BD. The presence of an incomplete or disrupted zNA-BD in E3L homologues highly suggests that these proteins cannot fully compete with ZBP1 to inhibit necroptosis induction. However, further studies will be necessary to fully comprehend the action of these proteins regarding complete necroptosis inhibition.

## Discussion

Necroptosis is an inflammatory form of cell death that is mediated by RIPK3 and MLKL and provides an extra defense mechanism against pathogen infection, facilitating the elimination of virus-infected cells before the production of progeny virions (Upton and Chan 2014; Nogusa et al. 2016; Xia et al. 2020). Given the crucial role of necroptosis in the innate immune response of humans and mice (Orzalli and Kagan 2017; Nailwal and Chan 2019), it was broadly accepted that this pathway was ubiquitous in mammals. Our results from 67 species across nine mammalian lineages provides the first comprehensive picture of the molecular evolution of necroptosis in mammals. We show that while RIPK1 is under evolutionary conservation, RIPK3 and MLKL are poorly conserved in lineages that evolved separately over the course of evolution.

A detailed analysis of RIPK3 and MLKL in mammals reveals a complex pattern where lagomorphs, cetaceans, carnivores and species from rodent and afrotheria lineages separately lost key components of the necroptotic pathway (Fig. 2A). The order lagomorpha includes two big families, Ochotonidae and Leporidae (Melo-Ferreira et al. 2015). The presence of the same frameshift mutation in Leporid species (+G, Fig. 3A) suggests that the disruption of the necroptotic pathway occurred early, but only after the bifurcation between Ochotona and Leporid given that the plateau Pika presents intact RIPK3 and MLKL proteins (Fig. 2A). Despite all efforts, and despite the complete genome assembly surrounding the MLKL flanking genes in the European rabbit genome, no partial or complete MLKL gene was found, indicating that this gene is deleted in the European rabbit genome and possibly in the remaining Leporid species (Fig. 4). The core genes of the necroptotic pathway also presented premature stop codons in cetaceans. In the studied cetaceans, the MLKL gene presented a common stop codon in the first exon, resulting in the inactivation of this gene (Fig. 3B). Again, the presence of similar patterns of pseudogenization in RIPK3 or MLKL genes within species of the same order infer that disruption of these genes occurred before their diversification and was maintained throughout evolution. On the other hand, *RIPK3* disruption in cetaceans appears to be the result of insertions or deletions that are not shared between closely related species, but rather specific to each species (S Appendix 6), suggesting that these disrupting mutations occurred later in evolution when compared to MLKL. The addition of some rodents as well as afrotheria species to the list of mammals that have disrupted necroptotic pathways, raises the possibility that other closely related species might have lost this pathway after the diversification of these lineages. It is currently believed that activation of the RIPK3 and recruitment of MLKL are critical steps during necroptosis. For example, deleting either RIPK3 or MLKL can lead to the suppression of skin and liver inflammation in mice (Dannappel et al. 2014; Rickard et al. 2014). Moreover, when mice are treated intravenously with a high-doses of TNF, there appears to be no differences between RIPK3-deficient and MLKL-deficient mice (Moerke et al. 2019), substantiating the premise that MLKL follows RIPK3 in the necroptotic signalling. However, this appears not to be the case for all necroptotic cell death responses, as different studies revealed alternative pathways for MLKL and RIPK3-dependent programmed necrosis that are executed in the absence of RIPK3 or MLKL, respectively (Günther et al. 2016; Zhang et al. 2016). To date, there are no studies suggesting that RIPK3/MLKL double-knockout mice are still able to induce necroptosis, which indicates that species that have disrupted RIPK3 and MLKL lost the necroptotic pathway throughout evolution.

The loss of function of *RIPK3* and *MLKL* in independently evolving lineages (convergent evolution) indicates that gene loss is an important evolutionary mechanism for phenotypic change in these animals and may contribute to similar adaptations. Even though it would be expected that loss of genes is maladaptive, gene loss can be beneficial by providing an evolutionary mechanism for adaptations. In fact, if the loss of an existing gene would increase fitness by making a species better adapted to the environment that surrounds it, then gene loss would be an easy solution to an evolutionary problem. Necroptosis contributes to innate immunity as a pathogen clearance mechanism (Xia et al. 2020). However, contrary to apoptosis, in which several highly immunogenic intracellular proteins are sequestered in the dead cell, necroptosis releases DAMPs in the surrounding tissue that promote strong inflammatory responses and result in the attraction of different types of immune cells to the site of infection (Kaczmarek et al. 2013). Studies in mouse models have provided strong evidence that necroptosis is implicated in several inflammatory neurodegenerative diseases, including multiple sclerosis and amyotrophic lateral sclerosis (Ofengeim et al. 2015; Ito et al. 2016). Mouse-model experiments identified keratinocyte necroptosis as a trigger of skin inflammation (Bonnet et al. 2011) and a correlation between necroptosis and intestinal inflammation has also been established (Welz et al. 2011; Pierdomenico et al. 2014). Thus, while necroptosis might mediate host defense, its inhibition in certain contexts may lessen disease severity. It is known that excessive inflammation can promote cancer cell growth and metastasis (Najafov et al. 2017). Thus, it is possible that a pro-inflammatory cell death like necroptosis might promote metastasis and thus, inhibition of this pathway might represent an advantage for regulation of cancer cell growth. Intriguingly, some of the species that are lacking the core necroptotic machinery are known to resist cancer. That is the case for cetaceans, the naked mole-rat and african elephants (Liang et al. 2010; Abegglen et al. 2015; Tejada-Martinez et al. 2021). It is also possible that selection against necroptosis in different mammalian lineages could have been driven by different factors depending on the environment or conditions. For example, it was previously suggested that the absence of MLKL in Carnivores reflected a microbe-rich and virus-containing diet of raw meat, causing evolutionary counter-selection against necroptosis (Dondelinger et al. 2016). Nevertheless, the absence of the necroptotic pathway in independently evolving lineages suggest that the deregulation of this pathway was detrimental for the host organism, which ultimately drove selection against the presence of RIPK3 and MLKL.

As many other viruses, poxviruses express immunomodulatory and host-range factors important for the suppression and evasion of the host innate and adaptive antiviral responses (Werden et al. 2008; Oliveira et al. 2017). VACV protein E3 not only sequester dsRNA through their dsRNA-BD limiting the activation of the innate immune system against the virus infection, but also inhibit the IFN-induced dsRNA dependent protein kinase (PKR), known to be a crucial component of the host innate immunity against viral infection, replication, and spread (Davies et al. 1993; Sharp et al. 1998). Our results show that the dsRNA-BD of distant E3L proteins present high levels of sequence similarity (Fig. 5B and C), which is consistent with the ability of this domain to target conserved pathways present in different hosts. Although the dsRNA sequestration functions of the E3 C-terminal have been clear for decades (Chang and Jacobs 1993; Thompson et al. 1994), the IFN sensitivity of VACV E3 N-terminal deletion mutants remained unresolved for a long time. Recently, strong evidence showed that the E3 N-terminal domain competes with ZBP1 to prevent ZBP1-dependent activation of RIPK3 and consequent necroptosis (Koehler et al. 2017; Koehler et al. 2020). The model proposed by the authors suggests that during WT-VACV infection, the zNA-BD of E3 binds to VACV-induced Z-form nucleic acid and masks it, preventing sensing by ZBP1 and further RIPK3 necroptosis induction (Koehler et al. 2017; Koehler et al. 2020). However, it is interesting that poxviruses like MPXV, MYXV and CePV-TA have E3L homologs that present a complete dsRNA-BD but not zNA-BD (Fig. 5B). In VACV-E3 Δ83N–infected cells (mutant lacking the first 83 aa corresponding to the zNA-BD), the absence of the zNA-BD facilitates ZBP1 to sense VACV-induced PAMPs and initiate necroptosis induction (Koehler et al. 2017). Therefore, it is expected that E3L homologues that lack N-terminal zNA domains, like CePV-TA-21 and M029, cannot prevent Z-form nucleic acid sensing triggering necroptosis induction and early abortion of viral replication. Like CePV-TA-20, F3 protein is also missing several amino acids in the N-terminal region and presents high conservation in areas that are known to bind to zNA (Fig. 5C). Nevertheless, F3 protein seems to have lost the ability to compete with ZBP1 and inhibit sensing (unpublished data). It was previously shown that the N-terminus of VACV E3 is necessary for IFN resistance in JC cells since VACV-E3Δ37N (mutant mimicking MPXV E3 zNA-BD) did not initiate DNA replication (Arndt et al. 2015). However, MPXV was able to replicate efficiently in the same cells, despite having a partial N-terminal zNA-BD, suggesting that the predicted binding to z-form nucleic acid was intragenic and downstream of z-NA sensing, rather than related to the ability of F3 zNA-BD to mask z-form nucleic acid (Arndt et al. 2015).

Interestingly, inactivation of necroptosis in Lagomorphs and Cetaceans seems to correlate with the absence of the E3L zNA-BD in their naturally infecting poxviruses, namely leporipoxviruses (MYXV and SFV) and cetaceanpoxviruses (CePV-TA), respectively. Monkeypox is a viral zoonosis endemic to central and western Africa areas where African rope squirrels and other rodents are likely reservoir hosts (Essbauer et al. 2010). Interestingly, the absence of a functional N-terminus in MPXV F3 protein also seems to correlate with the fact that some rodents appear incapable of undergoing necroptosis. Like MPXV, leporipoxviruses and CePV-TA pathogenesis are restricted to only certain species and have little or no pathogenesis capability in all others (Rahman et al. 2013; Arndt et al. 2015; Rodrigues et al. 2020). Infection of the same host over hundreds of years or even millennia may drive the evolution of each virus to rapidly evolve to a fitness peak in a given host environment. Previous niche-filling models (Holt 2009; Cooper et al. 2010; Simmonds et al. 2019) emphasize the role of host interactions in shaping virus evolution. According to these models, as hosts diversify and speciate over longer evolutionary periods, viral host factors that aim to counter the host antiviral functions are subject to continuous changes. Indeed, it is known that genes associated with host antiviral mechanisms present high evolutionary rates and are often under positive selection (Münk et al. 2012; van der Lee et al. 2017; Águeda-Pinto, Lemos de Matos, et al. 2019). Here, we suggest that during the evolution of these poxviruses, the loss of the zNA-BD did not present a disadvantage in the host organism; therefore, this trait was maintained, which reflects how these viruses adapt as their niche changed.

### Concluding remarks

The disruption of necroptosis in independently evolving lineages suggests a convergent evolutionary loss of this pathway, probably reflecting an important selective mechanism for phenotypic change. Interestingly, we also found a strong correlation between the disruption of necroptosis in leporids and cetaceans and the absence of the E3L zNA-BD (responsible for necroptosis inhibition) in their naturally infecting poxviruses as in the case of MYXV and CePV-TA, respectively. Overall, our study provides the first comprehensive picture of the molecular evolution of necroptosis in mammals, highlighting the importance of gene/pathway loss for the process of species adaptation and suggesting that it is a true pathogen-response pathway that is not required for normal mammalian development. Moreover, this study sheds some light on a co-evolutionary relationship between poxviruses and their hosts, emphasizing the role of host adaptation in shaping virus evolution.

## Materials and Methods

### Genomic approach to detect genes associated with the necroptosis pathway

To detect intact and inactivated genes, we first identified the key genes of the necroptotic pathway (i.e., *RIPK1*, *RIPK3* and *MLKL*) in the human (*H. sapiens*) and mouse (*Mus musculus*) reference genomes and looked for the presence of orthologues in existing genome sequence databases from 67 different species that belong to the 9 main mammalian orders/superorders: primates, rodents, lagomorpha, chiroptera, carnivora, perissodactyla, artiodactyla, cetacea, and afrotheria (S Appendix 1). We did not only search for the complete loss of exons or entire genes, but also searched for insertions and deletions that shift the reading frame, frame-preserving insertions that create a premature stop codon and substitutions that create an in-frame stop codon. The respective search methodology had been previously applied to the identification of different homologues of different annotated mammalian genomes (Sharma et al. 2018; Águeda-Pinto, Castro, et al. 2019). To further ensure that all gene loss events discussed in this study are real and not due to sequencing errors, we validated them either by sequencing of samples or by using a curated bioinformatic pipeline (see below).

### Amplification and sequencing of RIPK1 and RIPK3 nucleic acid sequences from Lagomorpha species

In contrast to the majority of mammalian orders, lagomorpha only presents three annotated genomes: the European rabbit (*Oryctolagus cuniculus*, accession # GCA_000003625.1), the American Pika genome (*Ochotona princeps*, accession # GCA_014633375.1) and the plateau pika (*O. curzoniae*, accession # GCA_017591425.1). Given the importance of lagomorphs for this study, samples from different lagomorpha species were used to obtain the nucleic coding sequence from RIPK1. For that, RNA was extracted from tissues of *O. cuniculus cuniculus*, *O. cuniculus algirus*, *Lepus americanus*, *europaeus*, *L. timidus*, *L. granatensis*, *Sylvilagus floridanus*, *S. bachmanis*, *O. princeps* and *O. collaris* samples, using the Qiagen DNeasy Blood & Tissue kit (Qiagen, USA) following manufacturer’s instructions. Synthesis of cDNA was achieved by using SuperScript III Reverse Transcriptase (Invitrogen, USA). Primers were designed according to the *RIPK1* transcript from *O. cuniculus* [Accession # XM_017350509.1] (Forward 5’-ATGTCTTTGGATGACATTAAAATG-3’ and Reverse 5’-CTACTTCTGGCTGAGCTGTATC-3’) and used to amplify the samples mentioned before. Phusion^®^ High-Fidelity DNA Polymerase (Finnzymes, Espoo, Finland) was used in the PCR amplification and the conditions included an initial denaturation (98°C for 3min), 35 cycles of denaturation (98°C for 30s), annealing (60°C for 15s) and extension (72°C for 30s) followed a final extension (72°C for 5 min).

From our initial search, the *RIPK3* gene was not annotated in the European rabbit. However, after mapping the location of RIPK3 based on its location in *H. sapiens*, *M. musculus* and *O. curzoniae*, we were able to identify a partial RIPK3 sequence in the European rabbit genome that presented an early stop codon. To exclude potential artifacts that can mimic real gene-inactivating mutations, a forward (5’-ATGTCTTCTGTCAAATTGTGG-3’) and a reverse (5’-ACTGCCTGCATCAGGATC-3’) primer were designed based on the parcial *RIPK3* sequence and were used to amplify the same region in the genomes from *O. cuniculus cuniculus*, *S. floridanus*, *L. americanus*, *L. europaeus* and *L. saxatilis*. For that, genomic DNA was extracted using the Qiagen DNeasy Blood & Tissue kit (Qiagen, USA) according to the manufacturer’s instructions. Phusion^®^ High-Fidelity DNA Polymerase (Finnzymes, Espoo, Finland) was used in the PCR amplification and the conditions included an initial denaturation (98°C for 3min), 9 cycles of denaturation (98°C for 30s), annealing (66°C for 15s) and extension (72°C for 30s) followed by more 25 cycles of denaturation (98°C for 30s), annealing (61°C for 15s) and extension (72°C for 30s) and a final extension (72°C for 5min).

Amplicons sequencing from RIPK1 and RIPK3 was performed with the ABI PRISM BigDye Terminator v3.1 Cycle Sequencing Kit and according to manufacturer’s protocol; reactions were cleaned with Sephadex™ (GE Healthcare Life Sciences, UK) and applied on an ABI PRISM 310 Genetic Analyser (Life Technologies, Applied Biosystems, Carlsbad, CA, USA). The obtained RIPK1 coding sequences and the partial RIPK3 sequences from the different Lagomorphs have been deposited in the GenBank database under the accession numbers that are shown in S Appendix 2. All samples were supplied by CIBIO/InBIO, Vairão, Portugal and used in previous studies (de Matos et al. 2011; Águeda-Pinto, Lemos de Matos, et al. 2019). No animals were captured, handled, or killed specifically for the purpose of this study.

### Detailed analysis on the cetacean genomes

Briefly, NCBI gene annotations for the gene orthologues of *MLKL* and *RIPK3* were initially screened via PseudoChecker (pseudochecker.ciimar.up.pt), which evaluates the coding condition of a gene (Ranwez et al. 2018; Alves et al. 2020). For each gene, a PseudoChecker analysis was run (default parameters), using the *Bos taurus* (cow) gene orthologue as a comparative input (NCBI Accession ID regarding cow MLKL: XM_002694707.6; RIPK3: XM_024997365.1), as well as the genomic sequences encompassing the putative ORF of the orthologous counterpart of each target species, directly exported from the NCBI genome browser. Through PseudoIndex, a built-in assistant metric, we quickly assessed the erosion status of the tested genes on a discrete scale ranging from 0 (coding) to 5 (pseudogenized) (Alves et al. 2020). Subsequent manual annotation was performed by importing the previously collected genomic sequences into Geneious Prime 2020 software (www.geneious.com) (Kearse et al. 2012) and determining each gene’s CDS using as reference cow’s *MLKL* and *RIPK3* orthologues sequences. In detail, per gene and species, using the built-in map to reference tool (highest sensitivity parameter selected), each (3’ and 5’ untranslated region-flanked) reference coding-exon was mapped against each target genomic sequence. Exons alignments were further screened for gene disruptive mutations, including in-frame premature stop codons, frameshift, and splice site mutations (any deviation from the consensus donor splice site GT/GC or the consensus acceptor splice site AG).

To inspect if the identified genetic lesions were not rendered as result of sequencing and/or genome assembly artifacts, we performed mutational validation (one per gene and species), resorting of raw genomic sequencing reads, retrieved from two independent genomic projects from the NCBI sequence read archive (SRA), when available. Explicitly, blastn searches were directed to the selected SRA projects, using the nucleotide sequence portion containing the selected mutation(s) as a query. The matching sequencing reads were downloaded into Geneious Prime 2020 (Kearse et al. 2012) software and mapped against the manually annotated mutation (highest sensibility parameter selected), confirming, or not, the presence of the identified mutation.

### Phylogenetic and molecular evolutionary analyses

The complete dataset of RIPK1, RIPK3 and MLKL proteins was aligned in BioEdit Sequence Alignment Editor using Clustal W (Thompson et al. 1994), followed by manual corrections when necessary. Amino acid alignments were then used to infer Maximum Likelihood (ML) phylogenetic trees using MEGA X (Kumar et al. 2018), with the substitution models JTT+G+F, JTT+G and HKY+G+I, respectively; determined using ProtTest (Darriba et al. 2011).

Given the fact that RIPK3 and MLKL proteins are highly divergent across the studied mammalian species, we decided not to perform any evolutionary analysis using these alignments. To look for signatures of natural selection operating in the RIPK1 alignment, we used HyPhy software implemented in the Datamonkey Web server (Pond and Frost 2005), to detect codons under selection: the Single Likelihood Ancestor Counting (SLAC) model, the Fixed Effect Likelihood (FEL) method (Kosakovsky Pond and Frost 2005), the Random Effect Likelihood, the Mixed Effects Model of Evolution (MEME) (Murrell et al. 2012) and Fast Unbiased Bayesian AppRoximation (FUBAR) (Murrell et al. 2013) methods were used. To avoid a high false positive rate, codons with p-values <0.05 for SLAC, FEL and MEME models and a posterior probability >0.95 for FUBAR were accepted as candidates for selection. For a more conservative approach, only residues identified as being under positive selection in three or more ML methods were considered.

### Analysis of VACV E3 homologues

VACV E3 homologues encoded by different poxviruses (=11) were retrieved from the NCBI database (https://www.ncbi.nlm.nih.gov/) and aligned in the BioEdit Sequence Alignment Editor using Clustal W (Thompson et al. 1994), followed by manual corrections when necessary. Amino acid alignments of the representative E3-like proteins were used to generate schematic diagrams using the COBALT program from the NCBI database.

## Acknowledgments

This work was supported by Foundation for Science and Technology (FCT), which supported the doctoral fellowship of A.Á.-P. (ref. SFRH/BD/128752/2017) and the investigator grant of P.J.E. (IF/00376/2015). This article is also a result of the project NORTE-01-0145-FEDER-000007, supported by Norte Portugal Regional Operational Programme (NORTE2020), under the PORTUGAL 2020 Partnership Agreement, through the European Regional Development Fund (ERDF). GM and MMR’s research is supported by the National Institute of Health (NIH) grants R01AI080607 and R01AI148302.

## Supporting information

**S. Appendix 1** Accession numbers for *RIPK1*, *RIPK3* and *MLKL* genes found in different mammalian lineages.

**S. Appendix 2** Accession numbers for *RIPK1*, *RIPK3* obtained from different Leporid samples.

**S. Appendix 3** Phylogenetic analysis for RIPK1, RIPK3 and MLKL proteins from different mammalian lineages.

**S. Appendix 4** Positive and negative selection analyses for RIPK1 protein.

**S. Appendix 5** RIPK3 and MLKL protein alignment from species from rodent and afrotheria lineages.

**S. Appendix 6** Tables identifying RIPK3 and MLKL mutations and premature stop codons in Cetacea order.

