## Supplementary material for "Convergent loss of the necroptosis pathway in disparate mammalian lineages shapes virus countermeasures": S appendix: S Appendix 1.pdf

**S Appendix 1.** Accession numbers for *RIPK1*, *RIPK3* and *MLKL* genes found in different mammalian lineages.

| Superorder/Order | Superfamily/Family | Species name | *RIPK1* | *RIPK3* | *MLKL* |
|---|---|---|---|---|---|
| Primates | Hominidae | *Homo sapiens* | NM 001354930.1 | NM_006871.4 | NM_152649.4 |
| | | *Pan troglodytes* | XM 016954824.1 | XM 001169864.3 | XM_016930166.2 |
| | | *Pan paniscus* | XM 003805555.3 | XM 003809099.1 | XM_024925971.1 |
| | Cercopithecidae | *Papio anubis* | XM 003896962.4 | XM 003901654.4 | XM_017953558.2 |
| | | *Macaca mulatta* | XM 001091986.3 | XM 001114079.3 | XM_015126622.1 |
| | | *M. fascicularis* | XM 015449962.1 | XM 005560982.2 | XM_015443138.1 |
| | | *Rhinopithecus roxellana* | XM 010361800.1 | XM 010363712.1 | XM_010382960.1 |
| | Cebidae | *Cebus capucinus* | XM 017497471.1 | XM 017520641.1 | XM_017530561.1 |
| | | *Aotus nancymaae* | XM 021672660.1 | XM 012468022.1 | XM_012437528.1 |
| | | *Saimiri boliviensis* | XM 010338093.1 | XM 003924268.2 | XM_010350629.1 |
| Rodents | Muroidea | *Rattus norvegenicus* | NM 001107350.1 | NM 139342.1 | XM_008772572.2 |
| | | *Mus musculus* | NM 001359997.1 | AF178953.1 | BC023755.1 |
| | Chinchillidae | *Chinchilla lanigera* | XM 005398899.2 | XM 013510393.1 | XM_013503862.1 |
| | Caviidae | *Cavia porcellus* | XM 013151465.1 | XM 003474285.4 | XM_013154995.2 |
| | Octodontidae | *Octodon degus* | XM_004628459.2 | STOP | STOP |
| | Bathyergidae | *Heterocephalus glaber* | XM_013073264.2 | STOP | STOP |
| | | *Fukomys damarensis* | XM_010622756.3 | STOP | STOP |
| | Sciuridae | *Ictidomys tridecemlineatus* | XM 021727876.1 | XM 013364463.2 | XM_013365168.2 |
| | | *Marmota marmota* | XM 015493957.1 | XM 015491924.1 | XM_015496076.1 |
| Lagomorpha | Ochotona | *Ochotona princeps* | XM_004596516.1 | INC | --- |
| | | *Ochotona curzoniae* | --- | XM_040969693.1 | XM_040982371.1 |
| | Leporidae | *Oryctolagus cuniculus* | XM_017350507.1 | STOP | --- |
| Chiroptera | Vespertilionidae | *Myotis brandtii* | XM 005869628.2 | XM 005859571.2 | XM_014536746.1 |
| | | *Eptesicus fuscus* | XM 028136859.1 | XM 008158442.2 | XM_028142801.1 |
| | Miniopteridae | *Miniopterus natalensis* | XM 016220367.1 | XM 016205787.1 | XM_016197763.1 |
| | Phyllostomidae | *Desmodus rotundus* | XM 024552320.1 | XM 024556248.1 | XM_024555929.1 |
| | Hipposidenidae | *Hipposideros armiger* | XM 019668516.1 | XM 019663032.1 | XM_019646402.1 |
| | | *Pteropus alecto* | XM 025044923.1 | XM 006913529.3 | XM_025044559.1 |
| | | *Rousettus aegyptiacus* | XM 016166677.1 | XM 016120560.1 | XM_016143898.1 |
| Carnivora | Mustelidae | *Mustela putorius* | XM 004753625.2 | XM 004755192.2 | --- |
| | | *Enhydra lutris* | XM 022517756.1 | XM 022523405.1 | --- |
| | Otoriidae | *Callorhinus ursinus* | XM 025891041.1 | XM 025870243.1 | --- |
| | | *Zalophus californianus* | XM 027602761.1 | XM 027569817.1 | --- |
| | Odobenidae | *Odobenus rosmarus* | XM 004408406.2 | XM 004402153.1 | --- |
| | Ursidae | *Ailuropoda melanoleuca* | XM 002924876.3 | XM 011227269.2 | --- |
| | | *Ursus maritimus* | XM 008692863.1 | XM 008708807.1 | --- |
| | Canidae | *Canis lupus* | XM 022414382.1 | XM 025442787.1 | --- |
| | | *Vulpes vulpes* | XM 025982629.1 | XM 026007627.1 | --- |
| | Felidae | *Acinonyx jubatus* | XM 027040185.1 | XM 015086216.1 | --- |
| | | *Felis catus* | XM 023253722.1 | XM 003987566.5 | --- |
| | | *Panthera pardus* | XM 019419890.1 | XM 019429283.1 | --- |
| Perissodactyla | Equidae | *Equus asinus* | XM 014867439.1 | XM 014868621.1 | XM_014850092.1 |
| | | *Equus caballus* | XM 023624365.1 | XM 005603254.3 | XM_005608429.3 |

| | | | | | |
|---|---|---|---|---|---|
| | | *Ceratotherium simum* | XM 004432163.2 | XM 004421921.2 | XM_014792683.1 |
| **Artiodactyla** | **Bovidae** | *Bos taurus* | NM 001035012.1 | XM 005211281.3 | XM_024978879.1 |
| | | *Bubalus bubalis* | XM 025264923.1 | XM 006061455.1 | XM_025268400.1 |
| | | *Pantholops hodgsonii* | XM 005971736.1 | XM 005985345.1 | XM_005980145.1 |
| | | *Capra hircus* | XM 018038999.1 | XM 013966825.2 | XM_013970972.2 |
| | **Suidae** | *Sus scrofa* | MG586799.1 | XM 001927424.4 | MG543991.1 |
| | **Camelidae** | *Camelus dromedarius* | XM 010978020 | XM 010995799.1 | XM_010987761.1 |
| | | *Camelus ferus* | XM 006175577.2 | XM 006172835.2 | XM_014554645.1 |
| | | *Vicugna pacos* | XM 006198664.2 | XM 006217294.2 | XM_015241915.1 |
| **Cetacea** | **Delphinidae** | *Orcinus orca* | XM 004281071.2 | **STOP** | **STOP** |
| | | *Lagenorhynchus obliquidens* | XM 027128236.1 | **STOP** | **STOP** |
| | | *Tursiops truncatus* | XM_019945096.2 | **STOP** | **STOP** |
| | | *Globicephala melas* | XM_030881534.1 | **STOP** | **STOP** |
| | **Phocoenidae** | *Neophocaena asiaeorientalis* | XM_024738443.1 | **STOP** | **STOP** |
| | | *Phocoena sinus* | XM_032650323.1 | **STOP** | **STOP** |
| | **Monodontidae** | *Monodon monoceros* | XM 007452108.1 | **STOP** | **STOP** |
| | | *Delphinapterus leucas* | XM_022586536.2 | **STOP** | **STOP** |
| | **Lipotidae** | *Lipotes vexillifer* | XM 028478546.1 | **STOP** | **STOP** |
| | **Physeteridae** | *Physeter catodon* | XM 028164463.1 | **STOP** | **STOP** |
| | **Balaenopteridae** | *Balaenoptera acutorostrata* | XM 004281071.2 | **STOP** | **STOP** |
| **Afrotheria** | **Elephantidae** | *Loxodonta africana* | XM_010597547.2 | **STOP** | --- |
| | **Trichechidae** | *Trichechus manatus* | XM_023741740.1 | **STOP** | **STOP** |
| | | *Elephantulus edwardii* | XM 006888715.1 | XM 006903266.1 | XM_006878861.1 |
| | | *Echinops telfairi* | XM 004711762.1 | XM 004698921.1 | XM_004704794.1 |
| | | *Chrysochloris asiatica* | XM 006870555.1 | XM 006835518.1 | **STOP** |
