## Supplementary material for "Convergent loss of the necroptosis pathway in disparate mammalian lineages shapes virus countermeasures": S appendix: S Appendix 3.pdf

**S Appendix 4.** Phylogenetic analysis for RIPK1, RIPK3 and MLKL proteins from different mammalian lineages

**S Appendix 4.1.** Phylogenetic analysis of RIPK1 proteins from different mammalian lineages. Amino acid sequences from different RIPK1 proteins were used to construct a Maximum Likelihood tree using MEGA X [34]. The support of the resulting nodes was estimated using 1000 bootstrap replicates. Bootstraps values are indicated on the branches.

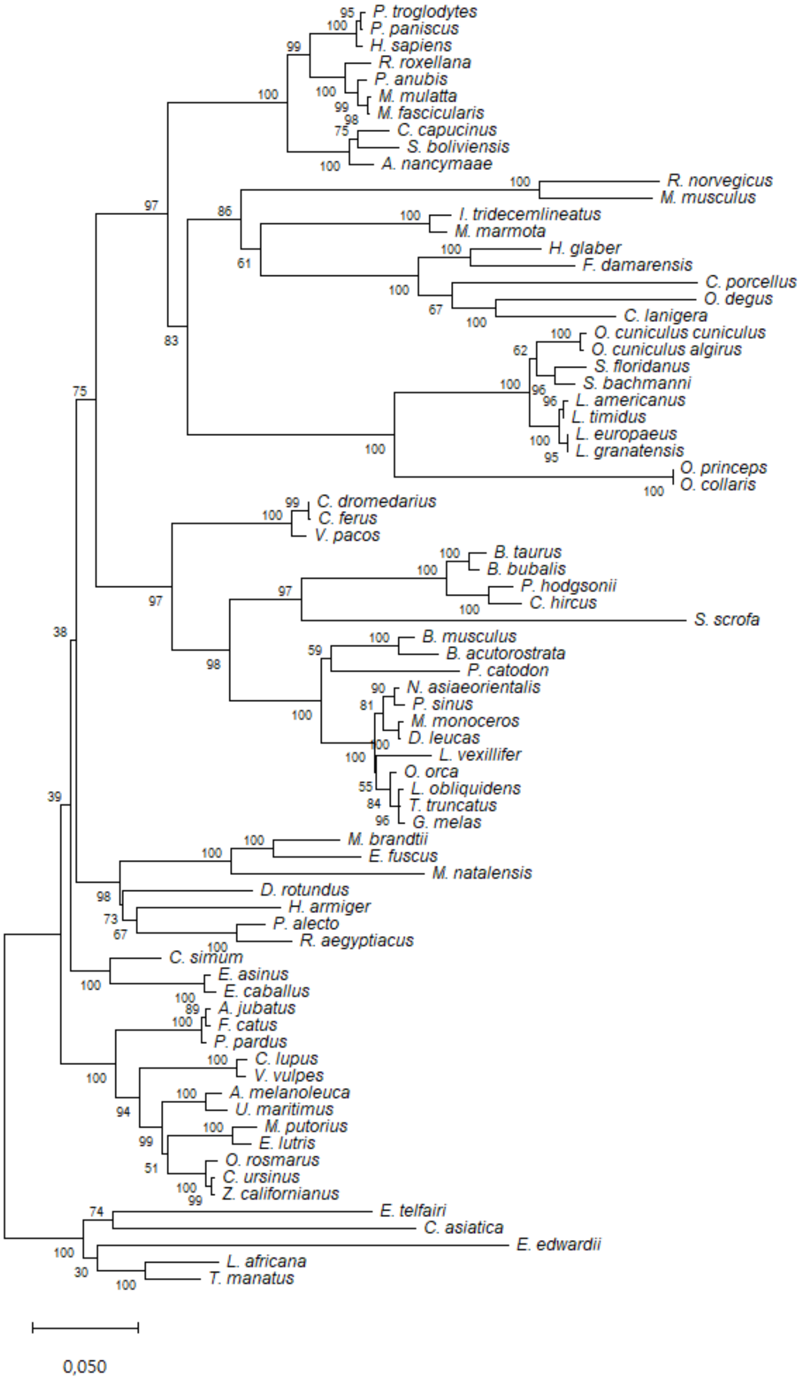

**S Appendix 4.2.** Phylogenetic analysis of RIPK3 proteins from different mammalian lineages. Amino acid sequences from different RIPK3 proteins were used to construct a Maximum Likelihood tree using MEGA X [34]. The support of the resulting nodes was estimated using 1000 bootstrap replicates. Bootstraps values are indicated on the branches.

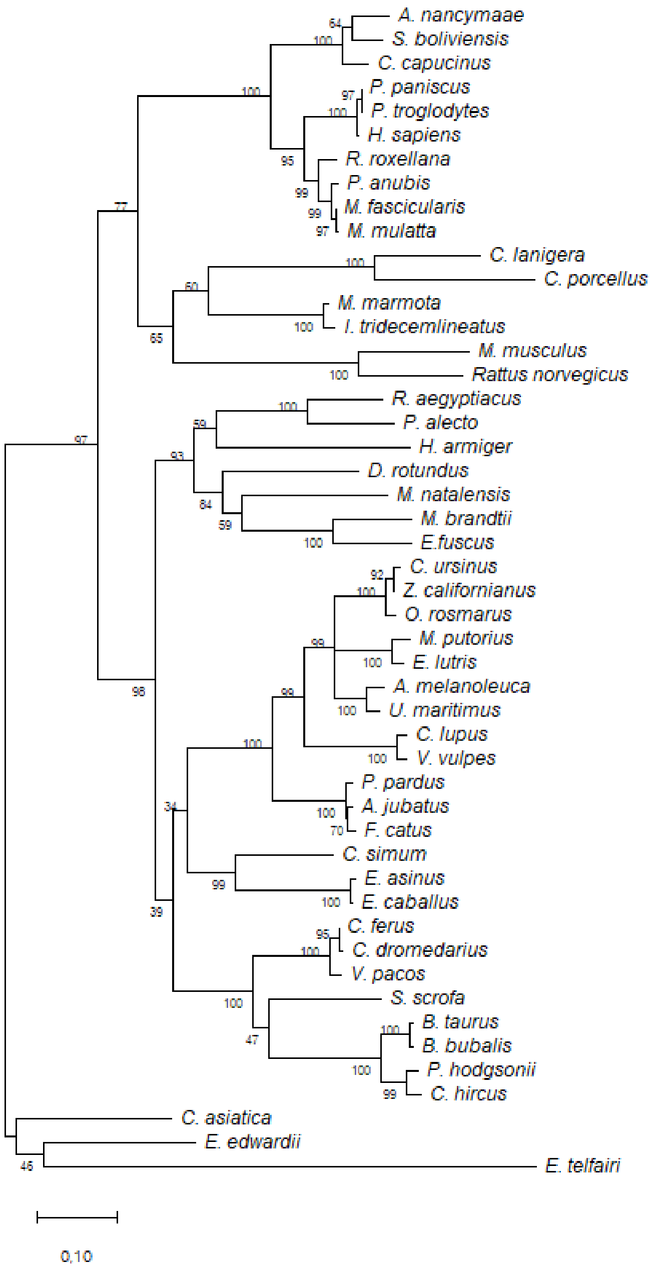

**S Appendix 4.3.** Phylogenetic analysis of MLKL proteins from different mammalian lineages. Amino acid sequences from different MLKL proteins were used to construct a Maximum Likelihood tree using MEGA X [34]. The support of the resulting nodes was estimated using 1000 bootstrap replicates. Bootstraps values are indicated on the branches.

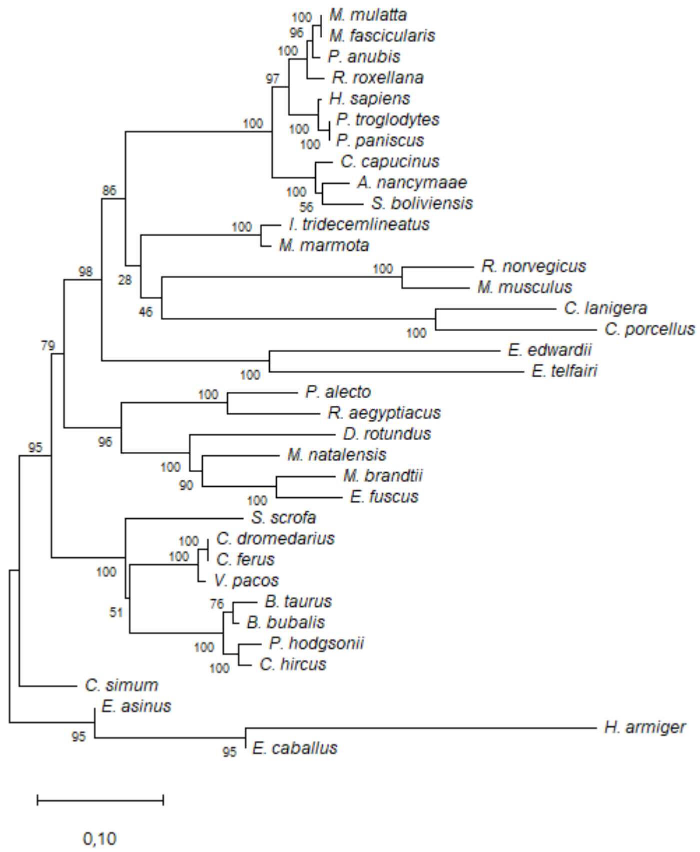
