## Supplementary material for "Convergent loss of the necroptosis pathway in disparate mammalian lineages shapes virus countermeasures": S appendix: S Appendix 4.pdf

**S Appendix 4.** Positive and negative selection analyses for RIPK1.

**S Appendix 4.1.** Positive selection using SLAC, FEL, MEME and FUBAR methods.

| Gene | Amino acids under Positive Selection in the alignment | | | | Total of sites* |
|---|---|---|---|---|---|
| | SLAC[a] | FEL[b] | MEME[b] | FUBAR[c] | |
| ***RIPK1*** | 183, 204, 258, 293, 294, 465, 501, 517, 518, 537, 539, 541, 543, 552, 689 | 183, 204, 258, 293, 294, 465, 501, 518, 537, 541, 543, 552, 597, 605, 689 | 19, 35, 54, 181, 184, 200, 204, 235, 236, 254, 265, 290, 293, 294, 315, 377, 490, 492, 518, 531, 537, 540, 541, 543, 546, 552, 579, 597, 626, 656, 679, 680, 679, 680, 681, 682, 683, 689, 691 | 204, 293, 294, 518, 543, 689 | **10** |

a)

Codons with significance level <0.1

b) Codons with significance level <0.05

c) Codons with posterior probabilities >0.90

*) Selected sites in three out of the four methods

**S Appendix 4.2.** Negative selection analysis using FEL method with negative residues selected at P > 0.5. Majority of negative selected residues fall in the first 300 aa and at the end of RIPK1 protein.

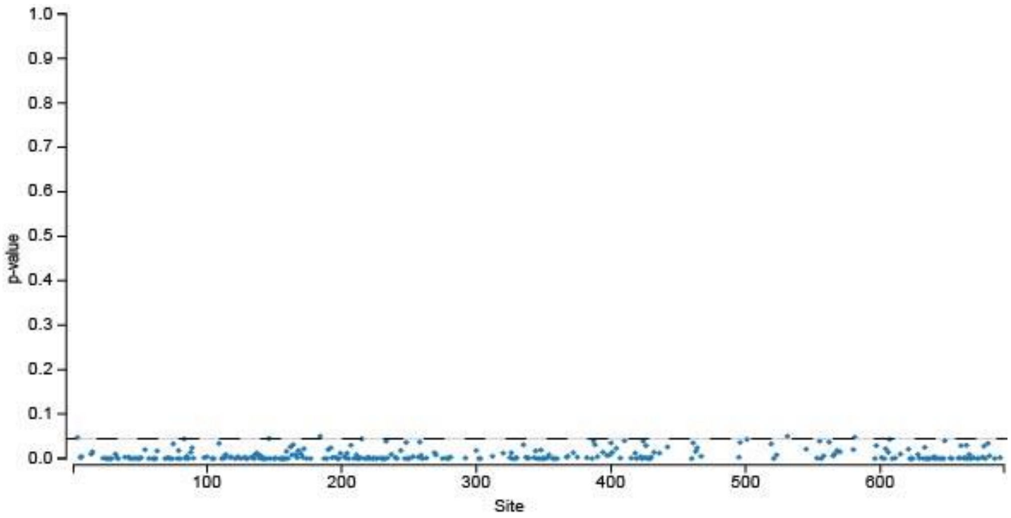

**S Appendix 4.3.** RIPK1 protein alignment used for positive and negative analysis.

```
                                    10        20        30        40        50        60        70        80        90       100
                           ....|....|....|....|....|....|....|....|....|....|....|....|....|....|....|....|....|....|....|....|
H. sapiens                 MQPDMSLNVIKMKSSDFLESAELDSGGFGKVSLCFHRTQGLMIMKTVYKGPNCIEHNEALLEEAKMMNRLRHSRVVKLLGVIIEEGKYSLVMEYMEKGNL
P. troglodytes             ....................................................................................................
P. paniscus                ....................................................................................................
P. anubis                  ........................................FV...........S................................N.............
M. mulatta                 ...................R....................FV......R....S................................N.............
M. fascicularis            ...................R....................FV......R.....................................N.........R...
R. roxellana               ----.....M.........R.....................V.........S..................................N.............
C. capucinus               .E..I...A....A.....R.................N...V......T..K.T..S..............................N.............
A. nancymaae               .E..L...A..........R.................N...V......T..K.T.................................N.............
S. boliviensis             .E..T...A..........R.................N...V......T..K.T.................................N.............
R. norvegicus              .......DN...A.D.L..RKD................H.FV.L.K..T...RA.Y.......G...H....D.......I.....N.........Q...
M. musculus                .......DN...A...L..KTD............Y..SH.FV.L.K..T...RA.Y..V....G...H............I.....N.............
H. glaber                  ----....D...N...L.QKED.......A....Y...H.FV.L.K..T..KRM.Y..S....G...H..S....MR.........N.............
F. damarensis              ----...DD...N...L.QKED.......A....Y...H.FV.L.K..T..KRM.Y..S....G...Y..S.R..M..........N.............
O. degus                   ----...DD.R.N..EL.WEED.A.....A.FS.Y...YEHV.L.K..T..KRM.Y.......G.V.HQ...RH..R...I.M...N.A....FV...S.
C. lanigera                ----...ED...N...L.WEED.......A......K.HKHV.L.K..T..KRM.Y..S....G...Y..S.N....................F......
C. porcellus               ----...DD...N.A.L.YKRN.......A.......MH..V.L.K..T..KRM.Y..S....G...R..S.........AV..D.N.C...........
I. tridecemlineatus        ----...DD...N...L.KKED................H..V.L.N..T..K.T.Y..S....G.I.H..K.........I.....D.............
M. marmota                 ----...DD...N...L.KKED................H.FV.L.K..T..K.T.Y..S....G.I.H..K.........I.....D.............
O. cuniculus cuniculus     ----...DD..........K.D.......E.......LH..V.L.K..T..KRA.Y..S....GR..H..K.....E....VM...N.C.........S.
O. cuniculus algirus       ----...DD..........K.D.......E.......LH..V.L.K..T..KRA.Y..S....GR..H..K.....E....VM...N.C.........S.
S. floridanus              ----...DD..........K.D.......E.......LH..V.L.K..T..KRA.Y..S....G...Y..K.....E...I.M...N.............
S. bachmanni               ----...DD..........K.D.......E.......LH..V.L.K..T..KRA.Y..S....G...Y..K.....E...I.M...N.............
L. americanus              ----...DD..........K.D.......E.......LH..V.L.R..T..KRA.Y..S....G...H..K.....E...I.M...N.............
L. timidus                 ----...DD..........K.D.......E.......LH..V.L.R..T..KRA.Y..S....G...H..K.....E...I.M...N.............
L. europaeus               ----...DD..........K.D.......E.......LH..V.L.R..T..KRA.Y..S....G...H..K.....E...I.M...N.............
L. granatensis             ----...DD..........K.D.......E.......LH..V.L.R..T..KRA.Y..S....G...H..K.....E...I.M...N.............
O. princeps                ----...DD.......L..KED...............LH..V.L.K..T..KRTDY..S....G...Y..K.....E.......D.N.............
O. collaris                ----...DD.......L..KED...............LH..V.L.K..T..KRTDY..S....G...Y..K.....E.......D.N.............
M. brandtii                ----...DD........Q.E.P...............SH..V.L.K..T..KRT.Y.AS....G...H....E.......I.M...N.............
E. fuscus                  ----...DD.........DTEH............L..SH..V.L.K..T..KRT.Y..S....G...H....E.......I.....N.............
M. natalensis              ----...ED.Q..PG.L..REH.......R....L...H..V.L.K..T..K.T.Y..S....G...H....E.......I.L...N...........S.
D. rotundus                ----...DD..........QEP.C.....Q....L..SY.FV.L.K..T..K.L.Y..S....G...H....E.......L.....N.C...........
H. armiger_                ----...DD.......V..KED............L..SH.FV.L.K..T..KRT.Y..S....G...H..S.Q.......I.....N.............
P. alecto                  ----...DD..........KEH............L..SH..V.L.K..T..KRT.Y..S....G...HK...Q.......I.....N.............
R. aegyptiacus             ----...DD..........KEF............L..SH..V.L.K..T..KRT.Y..S....G...HK...Q.......I.....N.............
M. putorius                ----...DD....T..L..KEY............L..CH..V.L.K..T..KRT.Y..S....G...H..................N.............
E. lutris                  ----...DD....T..L..KEY............L..CH..V.L.K..T..KRT.Y..S....G...H....N.............N.............
C. ursinus                 ----...DD..........KEH............L..SH..V.L.K..T..KRT.Y..S....G...H....N.........M...S.............
Z. californianus           ----...DD..........KEH............L..SH..V.L.K..T..KRT.Y..S....G...H....N.........M...S.............
O. rosmarus                ----...DD..........KEH............L..SH..V.L.K..T..KRT.Y..S....G...H....N.........M...S.............
A. melanoleuca             ----...DD..........KEH............L..SH..V.L.K..T..KRT.Y..S....G...H..................N......F......
U. maritimus               ----...DD..........KEH............L..SH..V.L.K..T..KRT.Y..S....G...H..................N.............
C. lupus                   ----...DD..........KEH............L..SH..V.L.K..T..KRT.Y..S....GR..H....R.........L...N.............
V. vulpes                  ----...DD..........KEH............L..SH..V.L.K..T..KRT.Y..S....GR..H....R.........L...N.............
A. jubatus                 ----...DD..........KEH............L..SH..V.L.K..T..K.T.Y..S....GR..H....N.........L...N.............
F. catus                   ----...DD..........KEH............L..SH..V.L.K..T..K.T.Y..S....GR..H....N.........L...N.............
P. pardus                  ----...DD..........KEH............L..SH..V.L.K..T..KRT.Y..S....GR..H....N.........L...N.............
E. asinus                  ----...DD.....T....QEY............LN.SHE.V.L.K..T..KRT.YD.S....GR..H....D.......I.....N.A......T....
E. caballus                ----...DD.....T....QEY............LN.SHE.V.L.K..T..KRT.YD.S....GR..H....D.......I.....N.A......TN...
C. simum                   ----...DD..........KEY............L..SH..V.L.K..T..KRT.Y..S....G...H....N.......I.....N.............
B. taurus                  ----...DD...S.T....Q.D.........C..L..SH..V.L.K..T..KRT.Y.......GR..H....R.........M...S.........Q.D.
B. bubalis                 ----...DD...S.T....Q.D.........C..L..SH..V.L.K..T..KRT.Y.......GR..H....H.........M...S.........Q.D.
P. hodgsonii               ----...DD...SPT....Q.D.........C..L..SH..V.L.K..T..KRT.Y.......GR..H....C.........M...S.........Q.D.
C. hircus                  ----...DD...SPK....Q.D.........C..L..SH..V.L.K..T..KRT.Y.......GR..H....C.........M...S.........Q.D.
S. scrofa                  ----...DD..........KED.E............KSH..V.L.K..T..KRN.Y..S....GR..H........R....VM...D....I...A..D.
C. dromedarius             ----...DD.....A....QED.......R....L...H..V.L.K..T..K.T.Y..S....GR..H........Q.....M...S.............
C. ferus                   ----...DD.....A....QED.......R....L...H..V.L.K..T..K.T----------------------------------------------
V. pacos                   ----...DD.....A....QED.......R....L...H..V.L.K..T..K.T.Y..S....GR..H........Q.....M...S.............
O. orca                    ----...DD.R.S.TE..RQED............L.SSHE.VVL.K..T..KRT.YD......GR..H..............M.D.N.......V...D.
L. obliquidens             ----...DD.R.S.TE..KQED............L.SSHE.VVL.K..T..KRT.YD......GR.RH..............M.D.N.......V...D.
T. truncatus               ----...DD.R.S.TE..KQED............L.SSHE.VVL.K..T..KRT.YD......GR.RH..............M.D.N.......V...D.
G. melas                   ----...DD.R.S.TE..KQED............L.SSHE.VVL.K..T..KRT.YD......GR.RH..............M.D.N.......V...D.
N. asiaeorientalis         ----...DD.R.S.TE..RQED............L.SSH..VVL.K..T..KRT.YD.....DGRT.H..............M.D.N.......V...D.
P. sinus                   ----...DD.R.N.AE..RRED............L.SSH..VVL.K..T..KRT.YD.....DGR..H..............M.D.N.......V...D.
M. monoceros               ----...DD.R.S.TE..RQED............L.SSH..VVL.K..T..KRT.YD.....DGR..H..............M.D.N.......V...D.
D. leucas                  ----...DD.R.S.TE..RQED............L.SSH..VVL.K..T..KRT.YD.....DGR..H..............M.D.N.......V...D.
L. vexillifer              ----...DD.R.S.TE..RQED..G.........L.SSH..VVL.K..T..KLT.YD......GR..H..............M.D.N.......V...D.
P. catodon                 ----...DD.E.N.T....REG............L..SH..VVL.K..T..KRT.Y.......GR.RH..............M...N.......V...D.
B. musculus                ----...DD...N.A...LQKD..G.........L..SH..VVL.K..T..RRA.YS......GRV.H........Q.....M.D.N.......L...D.
B. acutorostrata           ----...DD...N.A...QQKD..G.........L..SH..VVL.K..T..KRA.YS......GRV.H..Q.....Q.....M.D.N.......L...D.
L. africana                ----...DD.....T...QTTH............H..IH..V.L.N..T..K.T.Y..S....G...H..N.N...........D.N.............
T. manatus                 ----...DD.....T...QT.H............C..LH..V.L.N..T..KRT.Y..S....G...H..N.N...........D.N.............
E. edwardii                ----...DD.E...T.L.VKK.I.N.S..E....L...C..V.L.K..T..K.T.Y..S....G...H..N.N...........D.N.............
E. telfairi                ....VF.DE.....T...TTRYI..W..........KIHRNV.L.N..T..K...YD.S....G...H..N.H.........M...S........K....
C. asiatica                ....TC.DD..L..T...NAVH.........C..K.IIL.YV.L.K..T...RA.Y.......GR..H..N.D..M..........N.C......K....

                                   110       120       130       140       150       160       170       180       190       200
                           ....|....|....|....|....|....|....|....|....|....|....|....|....|....|....|....|....|....|....|....|
H. sapiens                 MHVLKAEMSTPLSVKGRIILEIIEGMCYLHGKGVIHKDLKPENILVDNDFHIKIADLGLASFKMWSKLNNEEHNELREV-DGTAKKN-GG-TLYYMAPEH
P. troglodytes             ...............................................................................-....R..-..-.........
P. paniscus                ...............................................................................-....R..-..-.........
```

| | |
|---|---|
| P. anubis | ...................V...........................D....................SK......K..-.S.S...-..-......... |
| M. mulatta | ...................V...........................D....................SK......K..-...S...-..-......... |
| M. fascicularis | ...................V...........................D....................SK......K..-...S...-..-......... |
| R. roxellana | .........I.........V...........................D....................SK......K..-..AS...-..-......... |
| C. capucinus | ...................V...........................D...............T....SK.........-...S...-..-......... |
| A. nancymaae | ...................V...........................D...............T....SK.........-.S.S...-..-......... |
| S. boliviensis | .......V...........V...........................D...............T....SK.........-.S.S...-..-......... |
| R. norvegicus | .....TKE.V.........V......H...DE...............R..........V....T....TK....KQ..A-SSVT...-..-......... |
| M. musculus | .....TQIDV...L.....V.A........D................R..........V....T....TK.KD.KQK..-SS.T...N..-......... |
| H. glaber | .Q..R.QI.I.........A...........A..........................V....T....TK.....Q.KA-NS.S...-..-......... |
| F. damarensis | .Q....Q..I.....A.V.A...........A..........................V....T....TK.....Q.KA-NS.S...-..-......... |
| O. degus | ..M...QI.I....R...VA.T.........E.........D.L...K..........V....T....TK.....Q.RA-N...RR.-..-N........ |
| C. lanigera | .N....QI.I.........A.T.........E..........................V....T....TK.....Q.KA-.S.T...-..-......... |
| C. porcellus | L...Q.QT-I.........V.T.........E...............S..........V....T..R.TQ.....Q.KA-S..TRN.-..-..H...... |
| I. tridecemlineatus | .Q....QI.I.........V.T.........E...............DN.........V....T....TK.K...Q.K.-NSNSV..-..-......... |
| M. marmota | .Q....QI.I.........M.T.........E...............DN.........V....T....TK.K...Q.KM-NSNSV..-..-......... |
| O. cuniculus cuniculus | .......I.I.........A.T.........E..........................V....T....TK.....Q.K.-KS.R..T-..-..C...... |
| O. cuniculus algirus | .......I.I.........A.T.........E..........................V....T....TK.....Q.K.-NS.R..T-..-..C...... |
| S. floridanus | .......I.I.........A.TV........E...............S..........V....T....TK.....Q.K.-NS.R..S-..-..C...... |
| S. bachmanni | .......I.I.........A.T.........E...............S..........V....T....TK.....Q.K.-NS.C..S-..-..C...... |
| L. americanus | .......I.I........VA.T.........E..........................V....T....TK.....Q.K.-NH.S...-..-......... |
| L. timidus | .......I.I........VA.T.........E..........................V....T....TK.....Q.K.-NH.S...-..-......... |
| L. europaeus | .......I.I.........A.T.........E..........................V....T....TK.....Q.K.-NR.S..S-..-......... |
| L. granatensis | .......I.I.........A.T.........E..........................V....T....TK.....Q.K.-NR.S..S-..-......... |
| O. princeps | .Q....KINI.........A.T.........E..........................V....T....TT.....Q.KA-HSAS...-..-......... |
| O. collaris | .Q....KINI.........A.T.........E..........................V....T....TT.....Q.KA-HSAS...-..-......... |
| M. brandtii | .......I.I.........M...........E...............DN.........V....T....TK.....Q.KA-.SLS...-..-......... |
| E. fuscus | .......I.I.........M.T........RE...............DN.........V....T....TK.....Q.KADSLSK.NG-.--......... |
| M. natalensis | .......I.I.........M.T.........ESI........................V....T....TKDQ...Q.RA-.SLS...-..-..H...... |
| D. rotundus | .......I.I...........T....R....E..........................V....T....TK.....Q.KM-NSLSN..-..-......... |
| H. armiger_ | .......I.I.........M.T........KE.I.............D..........V....T....TK.....Q.K.-NSPK.NG-.T--........ |
| P. alecto | .....T.S.I.........M.T....R.....N..............D..........V....T....TK..D.KQ.K.-NSSS...-..-......... |
| R. aegyptiacus | .....T.S.I.........M.T....R.....H..............D..........V....A....TK.....Q.K.-NSPS...-..G......... |
| M. putorius | .......V.I...L.....M...........Q.I.............D..........V.........TK.....Q.KL-NRVS...-..-..C...... |
| E. lutris | .......V.I...L.....M...........Q.I.............D..........V.........TK.....Q.KL-NRVS...-..-..C...... |
| C. ursinus | .......V.I.........M.T.........Q...............D..........V.........TK.....Q.KL-NSVS...-..-..C...... |
| Z. californianus | .......V.I.........M.T.........Q...............D..........V.........TK.....Q.KL-NSVS...-..-..C...... |
| O. rosmarus | .......V.I.........M.T.........Q...............D..........V.........TK.....Q.KL-NSVS...-..-..C...... |
| A. melanoleuca | .......V.I.........M.T.........Q...............D..........V.T.......TK.....Q.KL-NSVS...-..-..C...... |
| U. maritimus | .......V.I.........M.TV........Q...............D..........V.........TK.....Q.KL-NSVS...-..-..C...... |
| C. lupus | .......V.I.........M.T.........E...............D..........V....T....TK..I..Q.KL-NSVS...-..-..C...... |
| V. vulpes | .......V.I.........M.T.........E...............D..........V....T....TK..I..Q.KL-NSVS...-..-..C...... |
| A. jubatus | .......V.I.........M.T.........E...............D..........V....T....TK..N..Q.KMNNSAP...-..-..C...... |
| F. catus | .......I.I.........M.T.........E...............D..........V....T....TK..N..Q.KMNNSAP...-..-..C...... |
| P. pardus | .......I.I.........M.T.........E...............D..........V....T....TK..N..Q.KMNNSAPE..-..-..C...... |
| E. asinus | .R.....I.I.........M.T.........E...............G..........V.........TK..N..Q.KL-NS.S...-..-..C...... |
| E. caballus | .R.....I.I.........M.T.........E...............G..........V....T....TK.....Q.KL-NS.S...-..-..C...... |
| C. simum | .R.....I.I.........M.T.........E...............D..........V....T....TK.....R.K.-NR.S...-..-..C...... |
| B. taurus | .R....QV.I.........M.T....R....E...............S..........V....T....TK.....Q.KA-G.S.G.S-..-..H...... |
| B. bubalis | .R....QV.I.........M.T....R....E...............S..........V....T....TK.....Q.KA-G.S.G.S-..-..H...... |
| P. hodgsonii | .R....QV.I.......V.M.T....R....E...............S..........V....T....TK..R..Q.KA-G.G--.S-..-..H...... |
| C. hircus | .R....QV.I.......V.M.T....R....E...............S..........V....T....TK.....Q.KA-G.S.G.S-..-..H...... |
| S. scrofa | .R....QTPV........VM.T....R....E...............D..........V....T....TK.....Q.K.-TS.....-..-..H...... |
| C. dromedarius | .....TQI.I.....A...M.T.........E...............S..........V.........TK.....EKKA-SSASQ..-..-..H...... |
| C. ferus | -----------V...A...M.T.........E...............S..........V.........TK.....EKKA-SSASQ..-..-..H...... |
| V. pacos | ......QI.I.........M.T.........E...............S..........V.........TK.....EKKA-SS.SQ..-..-..H...... |
| O. orca | .RA...QI.I.........M.T....R....E...............G.............L.T..E.MK..LTAQ.RG-NS.T.N--..-..H...... |
| L. obliquidens | .RA...QI.I.........M.T....R....E...............G.............L.T..E.MK..LTAQ.RG-NS.T.N--..-..H...... |
| T. truncatus | .RA...QI.I.........M.T....R....E...............G.............L.T..E.MR..LTAQ.RG-NS.T.N--..-..H...... |
| G. melas | .RA...QI.I.........M.T....R....E...............G.............L.T..E.MK..LTAQ.RG-NS.T.N--..-..H...... |
| N. asiaeorientalis | .RA...QI.I.........M.T....R....E.............I.G.............L.T..E.MK..LTAQ.RG-NS.T.N--..-..H.V.... |
| P. sinus | .RA...QI.I.........M.T....R....E.............I.G.............L.T..E.MK..LTAQ.RG-NS.T.N--..-..H...... |
| M. monoceros | .RA...QI.I.........M.T....R....E.............I.G.............L.T..E.MK..LTAQ.RG-NS.T.N--..-..H...... |
| D. leucas | .RA...QI.I.........M.T....R....E.............I.G.............L.T..E.MK..LTAQ.RG-NS.T.N--..-..H...... |
| L. vexillifer | .RT...QI.I.........M.T....R....E...............G.............L.T..E.MK..LTAQ.RG-KS.T.N--.S-..H.....C |
| P. catodon | T.....QI.I.........M.T....R...KE.............L.G..........F..L.T.NE.MK..LTAQ.RG-NSST.S--..-..H...... |
| B. musculus | ......QI.I.........M.T........RE.............M.G...V......F..L.T..E.MR..LTAQ.RG-NS.T.D--..-S.H.T.... |
| B. acutorostrata | ......QI.I.........M.T........RE.............M.G..........F..L.T..E.MR..LAAQ.R--GNSTT.D-S.-S.H.T.... |
| L. africana | .Q...G.V.I...........T.........E.L.............D..........V....T....TK.....Q.K.-NC.S.R.-..-......... |
| T. manatus | .Q...G.V.I...........TV........E...............D..........V....T....TK.....Q.K.-NC.S.R.-..-..C...... |
| E. edwardii | .R.....I.I...........T........RE.I..............Y.........V.T..T....TK.....Q.KQ-TFISRR.-A.-..C...... |
| E. telfairi | .Q...W.I.I...........T.........E...............D..........V.........TK..R..Q.KM-NSAS.SS-..-..C...... |
| C. asiatica | .Q..TDKVRI...........T.........A...............D..........V....T....TK....LQ.KM-TTPS...-..-.VC.I.... |

| | |
|---|---|
| | 210 220 230 240 250 260 270 280 290 300 |
| | ....\|....\|....\|....\|....\|....\|....\|....\|....\|....\|....\|....\|....\|....\|....\|....\|....\|....\|....\|....\| |
| H. sapiens | LNDVNAKPTEKSDVYSFAVVLWAIFANKEPYENAICEQQLIMCIKSGNRPDVDDITEYCPREIISLMKLCWEANPEARPTFPGIEEKFRPFYLSQLEESV |
| P. troglodytes | .............................................................................................N...... |
| P. paniscus | .................................................................................................... |
| P. anubis | .....................................................A.I.H.......................................... |
| M. mulatta | .......................................................I............................................ |
| M. fascicularis | .......................................................I............................................ |
| R. roxellana | .........................................V...........E.I.................D.......................... |
| C. capucinus | ......R...........I...V...H.................I..........I..................L.V.........T............. |
| A. nancymaae | ...I..............I...V............................L...I.H.........Q......L.V.........E............. |
| S. boliviensis | ......R...........I...V................................I..................L.V.....................R. |
| R. norvegicus | .T.I.T............I..............V..TE.FLV..N.....N.E..L.F.........ER..QT...D....F...KE.K......F..Y. |
| M. musculus | ...I.............GI.......K......V..TE.FVI........N.EE.L...........ER..Q.I..D....L....E.......HF..Y. |

```
H. glaber                 .S.I.VR..........GI................SAEH.LI..RT....N.EE...R..K...H..E....ED..V.....S...Q.K...E.E...F.
F. damarensis             .CNI.VR..........GI................SAEH.LI..RN....N.EE.I.H..K...H..E....E...V..V..S...Q.K...E.EF....
O. degus                  .SNI.I...........GI..............T..VE..LL...G....N..E...R..KGV.D......RED..L.......D.Q.K.I...EF..F.
C. lanigera               .SNI.V...........GI..............T..VE...V...R.D...M.E.I.H..K...D..Q...KEE..L.....D....YK...V.E...F.
C. porcellus              .C.I.V...........GIM....I.D.....Y.LSAEH.LV...R.......K.I.H..T......TC..QED..L.........Q.K...E..F..F.
I. tridecemlineatus       ...I.............GI.................AE..LI........N.E..I.H..K......E........V.....S.........V..F..Y.
M. marmota                ...I.............GI.................AE..LI........N.E..I.H..K......E....T...V.....S.........V..F..N.
O. cuniculus cuniculus    ..................I....VL................L............VA.D..G.L.R..E.....D.R......D...R.....S.HY..F.
O. cuniculus algirus      ..................I....VL................L............VA.D..G.L.R..E.....D.RV.....D...R.....S.HY..F.
S. floridanus             ..................I......................L.............A.D..G.VVN..E...D.D.RV.....D...R.....S.HY..Y.
S. bachmanni              ..................I......................L.............A.D..G.VVN..E.....D.RV.....D...R.....S.HY..Y.
L. americanus             ...I..............I......................L........N....A.D..G.VVG..E.....D.R......D.........S.HY..Y.
L. timidus                ...I..............I......................L........N....A.D..G.VVG..E.....D.R......D.........S.HY..Y.
L. europaeus              ...I..............I......................L........N....A.D..G.VVG..E.....D.R...M..D.........S.HY..Y.
L. granatensis            ...I..............I......................L........N....A.D..G.VVG..E.....D.R...M..D.........S.HY..Y.
O. princeps               ..................I......................L...........C.I.H......G..Q...NED.R......D..GN.K...S.N...N.
O. collaris               ..................I......................L...........C.I.H......G..Q...NED.R......D..GN.K...S.N...N.
M. brandtii               ...I..R.S.........I................N....LI........N.N..I....K.......Q....D..V....R....Q......HN...DI
E. fuscus                 ...I..R.S.........I......................I........H....I.F..K.......Q....D.GV.........I......HN..DD.
M. natalensis             .DN...R.SD........I.......H..............I...A....N.N..I....K.......Q..DPD.QV....A...........HN...C.
D. rotundus               ...I..R.S.........I................T....V..........EG..I....K.......Q..V.D..L....A...........H..K.D.
H. armiger_               ...I..R.S.........I...............................N.E..I....K....I..E...T...V........D......S.....F.
P. alecto                 ...I..R.S.........I.......H.........................E..I.N..K...NI..Q.......V....A....N...........N.
R. aegyptiacus            ...I..R.S.........I.......H.........................E..I.N..K...NI..Q.......V....A....E...........N.
M. putorius               ...I....S........GI..................E...I........N.E..L....E....I..Q....K.DI....I.L........VD....N.
E. lutris                 ...I....S........GI..................E...I........N.E..L....E....I..Q....K.DI....T.L........VD....N.
C. ursinus                ...I....S........GI...............................N....L....V....I..Q..DTK..V....T..........VV....N.
Z. californianus          ...I....S........GI...............................N....L....V....I..Q..DTK..V....T..........VV....N.
O. rosmarus               ...I....S........GI...............................N....L....V....I..Q..DMK..V....T..........VV....N.
A. melanoleuca            ...I.T..S........GI...............................N.E..L....E....I..Q....K.......T..........VD....N.
U. maritimus              ...I....S........GI...............................N.G..L....E....I..Q....R.......T..........VD....N.
C. lupus                  ...I....S........GI......................L..........E..I....E....I..Q....K..I....A..........ED....N.
V. vulpes                 ...I....S........GI......................L..........E..I....E...GI..Q....K..I....A..........ED....N.
A. jubatus                ...I....S........GI......................I........N.E..I....E....I..Q....D..V....A..........VD....N.
F. catus                  ...I....S........GI......................I........N.E..I....E....I..Q....D..V....A..........VD....N.
P. pardus                 ...I....S........GI......................I........N.E..I....E....I..Q....D..V....A..........VD....N.
E. asinus                 ........S.........I......................I........N.E..I....K.......H.......V....A....N......D....N.
E. caballus               ........S.........I......................I........N.E..I....K.......H.......V....A....N......D....N.
C. simum                  ........S.........I.................................E..I....K.......Q.......V....V....N.....VN....N.
B. taurus                 .....SR.S.....F...I......................L..........E..I.F....V.DI.RQ...V..DN....A..........SN...DY.
B. bubalis                .....SR.S.....F...I......................L..........E..I.F....V.DI.RQ...G..DN....A.......H..SN...DY.
P. hodgsonii              .....SR.S.....F...I......................L..........E..I.F....V.D..RQ...V..DD....V..........SN...NY.
C. hircus                 .....SR.S.....F...I......................L..........E..I.F....V.D..RQ...V..DD....A..........SN...NF.
S. scrofa                 ......R.S.........I......................L........S....V.F....VVG...Q..ALS.DD....A.....Y.........GN.
C. dromedarius            ........S.........I......................V........S.G..I.H....V.GI..Q..DV...D....A..........SN....N.
C. ferus                  ........S.........I......................V........S.G..I.H....V.GI..Q..DV...D....A..........SN....N.
V. pacos                  ........S.........I......................V........S.G..I.H....V.GI..Q..DV...D....A..........SN....N.
O. orca                   ....SSR.S.....F...I....V..........V.......R....S....E...GH....AV.I.EQ..AV...D....A...........N....H.
L. obliquidens            ....SSR.S.....F...I....V..........V.......R....S....E...GH..W.AV.I.EQ..AV...D....A...........N....H.
T. truncatus              ....SSR.S.....F...I....V..........V.......R....S....E...GH....AV.I.EQ..AV...D....A...........N....H.
G. melas                  ....SSR.S.....F...I....V..........V.......R....S....E...GH....AV.I.EQ..AV...D....A...........N....H.
N. asiaeorientalis        ....SSR.S.....F...I....V..........P.......Y....S...TE...GH....AV.I..Q..AV...D....A...........N....H.
P. sinus                  ....SSR.S.....F...I....V..........P.......Y....S...ME...GH....AV.I..Q..AV...D....A...........N....H.
M. monoceros              ....SSR.S.....F...I....V..........P......L....G..G.E...GQ....AV.I..Q..AV...D....A...........N....H.
D. leucas                 ....SSR.S.....F...I....V..........P......L....G....E...GQ....AV.I..Q..AV...D....AC..........N....H.
L. vexillifer             ....SSR.S.....F...I....V..................R....S....E...GH....AV.I.EQ..AV...D....A....R......N....H.
P. catodon                ...A.SR.S.....F...M....V........G.......T.H.......A.E.V..Q..Q.A..I..Q..AV...D....A...........N....H.
B. musculus               ....SSR.S.....F...I....................V..RV...D....E.LA.H....A..I..Q..AVD.DD....A......K....N...GH.
B. acutorostrata          ....SSR.S.....F...I....V..................RV...D....E.LA.H....A..I..Q..AVH.DD....A......K....N....H.
L. africana               ..................I......................L........N.E..I.H..E.......H....S..V....A....NYK....K...KN.
T. manatus                ..................I......................I........N.E..I.H..K.......H.......V....A....N.KD...E...KD.
E. edwardii               .D.I..............I.......H.......MN.....A...N....H.N..I.H..K......RQ...SK.....P..A...T.KS.......DK.
E. telfairi               .....V............I......................I..T.....K.E..I....K...A...Q..A....V....A....N..S...KE...N.
C. asiatica               .S................I.............H........I........S....VK...Q...D...H............A....D.KS...KE...N.

                                   310       320       330       340       350       360       370       380       390       400
                          ....|....|....|....|....|....|....|....|....|....|....|....|....|....|....|....|....|....|....|....|
H. sapiens                EEDVKSLKKEYSN--ENAVVKRMQSLQLDCVAVPSSRSNSATEQPGSLHSSQGLGMGPVEESW---FAPSLEHPQ-EENEPSLQSKLQDEANYHLYGSRM
P. troglodytes            .............--................................................---.........-........................
P. paniscus               .............--................................................---.........-........................
P. anubis                 .............--Q..................P............................---.......Q.-............E...........
M. mulatta                .............--Q...............P..P............................---.......Q.-............E...........
M. fascicularis           .............--Q...............P..P............................---.......Q.-............E...........
R. roxellana              .............--Q..................P............................---.......R.-............E...........
C. capucinus              .............--.................L.P..................F.I.......---.......Q.-...V....N...E.T.........
A. nancymaae              .............--.................L.P..................F.I.......---....V..Q.-...V....N...E.T.........
S. boliviensis            .............--.................L.P..................F.V.......---.......Q.-...V....N...E.T.........
R. norvegicus             ....A......PS--QSP.L...F...H...PL.P.....--............P........---.SS.P.Y..-....R.V.A...E..S..AF.IFA
M. musculus               ....A......PD--QSP.LQ..F...H...PL.P.....--............Q........---.SS.P.Y..-D..DR.V.A...E..S..AF.IFA
H. glaber                 .K.L.......PM--Q.E.LR...........E.P.....--............EI.......---.SA....Q.-..H..I..H...E.....VF....
F. damarensis             .K.........PM--Q.E.FR.....R.....D.P.....--............E........---.SA.P.--.E..H.SN..C...E.....I.....
O. degus                  .K..E....D.PI--QKE.LR.RH...P...PE.P.....--............EI......D---.SA....Q.-..Y..I..H...K..D..AF.R..
C. lanigera               .K......E..PI--Q.EILR.VP.V.P...PE.P.....--...........PE........---.SA....Q.-..Y..I..R...K.....VF....
C. porcellus              .K.......K.PT--K.EALM.A........PE.L.....--..........P..T......Y---.SA.M.RQ.-..Y..I..H...E......F...I
I. tridecemlineatus       ....E......PM--K...LQ.L.........E.P.....--............QI.......---.SA...N..-..S..I..L...E..S..AF.N..
M. marmota                ....E......PM--K...LQ.L.........E.P.....--............QI.......---.S....N..-.DS..I..L...E..S..AF.N..
O. cuniculus cuniculus    .........R.PE--QS..............P..P..........S.......FMT....V.F---.SACP.IS.-............E...........
O. cuniculus algirus      .........R.PE--QS..............P..P..........S.......FMT....V.F---.SACP.IS.-............E...........
```

```
S. floridanus           .........R.PE--QS..............P..P..........S.......FMT....V.F---.SACP.IS.-............E...........
S. bachmanni            .........R.PE--QS..............P..P..........S.......FMT....V.F---.SACP.IS.-............E...........
L. americanus           ........RQ.PE--QS..............PA.P..........S.......FMT....V.F---.SACP.IS.-............E...........
L. timidus              ........RQ.PE--QS..............PA.P..........S.......FMT....V.F---.SACP.IS.-............E...........
L. europaeus            ........RQ.PE--QS..............PA.P..........S.......FMT....V.F---.SACP.IS.-............E...........
L. granatensis          ........RQ.PE--QS..............PA.P..........S.......FMT....V.F---.SACP.IS.-............E...........
O. princeps             .........D.PE--V...............PA.A..........N.......FQT....MTFASL.SA.P..L.-...Q........E...........
O. collaris             .........D.PE--V...............PA.A..........N.......FQT....MTFASL.SA.P..L.-...Q........E...........
M. brandtii             ....Q......PS--P.E.........I....I.P..........R......S.RI.......---....PDYQ.-....R...N...E...........
E. fuscus               ...........PS--Q.E.........I......P......ID..S........R........---....PDYQP-....Y...N...E......F....
M. natalensis           ..........FPS--Q.E.........I....I.P..........S........RI.......---....PD.Q.-.QS.LI..N...E......F....
D. rotundus             ...........PG--Q.EI........M....I.P..........C.................---....P..Q.-...DLR......E......F.N..
H. armiger_             ...........PC--Q.EM........M....I.P............................---.D..P..Q.-........N..........F....
P. alecto               ...........PS--Q.EI........M....I.P..........S........RI.......---....P..Q.-....L.......E......F....
R. aegyptiacus          ...........PS--Q.EI........M....I.P..........S........RI.......---....P..Q.-....L.......E......F....
M. putorius             .D..T....Q.PDPDQSSF....H...I..T.I.P......P...S.........I.......---......NQ.-...QL.......E......H....
E. lutris               .D..T......PDPVQ.SF....H...I..T.I.P......P...S.........I.......---......NQ.-...QL.......E......H....
C. ursinus              .D..T......PDPTQ.SF....H...I..T.I.P......P...S.....H...I.......---.S....NQ.-...QL.......E......F....
Z. californianus        .D..T......PDPDQ.SF....H...I..T.I.P......P...S.....H...I.......---.S....NQ.-...QL.......E......F....
O. rosmarus             .D..T......PDPAQ.SF....H...I..T....P......P...S.....H...I.......---.S....NQ.-...QL.......E......F....
A. melanoleuca          .D..T......PDAAQ.SF....H...I..T.I.P......P...S.........I.......---......NQ.-...QL.......E......F....
U. maritimus            .D..I......PDAAQ.SF....H...I..T.I.P......P...S.........I.......---.....DNQ.-...QL.......E......F....
C. lupus                ....T......PDPAQ.S.........I....T.P......P...S.........I.......---......NQ.-D..QL.......E...........
V. vulpes               ....T......PDPAQ.S.........I....T.P......P...S.........I.......---......NQ.-D..QL.......E...........
A. jubatus              ...........PD--Q.SF....H...I..I.I.P......P...S.................---......YQ.-....L..R....E...........
F. catus                ....T......PD--Q.SF....H...I..I.I.P......P...S.................---....V.YQ.-....L..R....E...........
P. pardus               ....T......PD--Q.SF....H...I..I.I.P......P...S.................---....V.YQ.-....L..R....E...........
E. asinus               ...........PD--QS.I........I....I.P..........S.................---...AS..Q.-....L.......E...........
E. caballus             ...........PD--QS.L........I....I.P..........S.................---...AS..Q.-....L.......E...........
C. simum                .D.........PG--QS.I........I....I.P..........S.................---....S..Q.-....LI......E......F....
B. taurus               ..........FPG--Q.EI....K...I....IAP..........S.......F.........---...APD.QP-...DFL.H....E...........
B. bubalis              ..........FPG--Q.EI....K...I....IAP..........S.......F.........---...APD.QP-...DFL.H....E...........
P. hodgsonii            .....R....FPG--Q.EI....K...I....IAP..........S.......F.........---...APD.QP-...DFL.N....E...........
C. hircus               ..........FPG--Q.EI....K...I....IAP..........S.......F.........---...APD.QP-...DFL.N....E...........
S. scrofa               .....N.....PS--Q...........I.....AP......P...........F.........---....A..Q-H....LH......E......F....
C. dromedarius          ...........PG--Q..I........I......P..........S.......FRI.......---.......E.-....L.......E...........
C. ferus                ...........PG--Q..I........I......P..........S.......FRI.......---.......E.-....L.......E...........
V. pacos                ...........PG--Q..I........I......P..........S.........I.......---.....A.E.-D...L.......E...........
O. orca                 ...........LG--Q.EI....K...IG...MSP..........S.....R.F.L....D..---...AP.QQ.QQ.EDLL.H....E...........
L. obliquidens          ...........LG--Q.EI....K...IG....SP..........S.....R.F.L....D..---....P.QQ.QQ.EDLL.H....E...........
T. truncatus            ...........LG--Q.EI....K...IG....SP..........S.....R.F.L....D..---....P.QQ---.EDLL.H....E...........
G. melas                ...........LG--Q.EI....K...IG....SP..........S.....R.F.L....D..---....P.QQ.Q..EDLL.H....E...........
N. asiaeorientalis      ...........LG--Q.EI....K...I....MSP..........S.......F.L....D..---...TP.QQ.QQ.EDLL.H....E...........
P. sinus                ...........LG--Q.EI....K...I....MSP..........S.......F.L....D..---...TP.QQ.QQ.EDLL.H....E...........
M. monoceros            ...........LG--Q.EI....K...IG...MSP......A...S.......F.L....D..---...AP.QQ.QQ.EDLL.H....E...........
D. leucas               ...........LG--Q.EI....K...IG...MSP......A...S.......F.L....D..---...AP.QQ.QQ.EDLL.H....E...........
L. vexillifer           ...........PG--Q.EI....K...I..I.ISP..........S.......F.L....D..---...AP.QR.QQQEDLL.H....E...........
P. catodon              ...........PG--Q.EI....K...I....ISP................R.F.S....D..---...AP.QQREWQEDVR.H....E..R........
B. musculus             ...........PG--Q.EI....K...I....I.P..........S.....R.F.L....D..---...AP.QQ.Q---DLR.H....E...........
B. acutorostrata        ...........PG--Q.EI....K...I....I.P..........S.....R.F.L....D..---...AP.QQ.Q---DLR.H....E...........
L. africana             ........D..LD--Q.VII.K.....T......P..........S........R........---.G..S..Q.-...Q...............F....
T. manatus              ........E..PD--Q.PII.K.....I....L.P..........S........R...I....---.G..S.YQ.-...Q........E......F....
E. edwardii             ........EA.PD--QDSFI.E.....I......P..................FR........---.G..S..Q.-.YE-L...N...E.....HF....
E. telfairi             DA.......WNLD--Q...F..L....I..T.I.P.....------------------------------------------------------------
C. asiatica             ...........PD--....I.......I..E.I.P..........S.......FR........---.G..S..Q.-....S......LE......F....

                                 410       420       430       440       450       460       470       480       490       500
                        ....|....|....|....|....|....|....|....|....|....|....|....|....|....|....|....|....|....|....|....|
H. sapiens              DRQTKQQPRQNVAYNREEERRRRVSHDPFAQQRPYENFQNTEGKGTAYSSAASHGNAVHQPSGLTSQPQVLYQNNGLYS--------SHGFGTRPLDPGT
P. troglodytes          ........................................M......................................--------.............
P. paniscus             ...............................................................................--------.............
P. anubis               ...M......................................V...............P.....S.....P.R..V...--------.............
M. mulatta              ...M......................................V...............P.....S.....P.R..V...--------.............
M. fascicularis         ...M......................................V...............P.....S.....P.R..V...--------.............
R. roxellana            ...M.....................................KV.........T.................P.R..V...--------.............
C. capucinus            ..........S.........K.........H...R..........I...G.T......Y........L..P.....S.G--------LR...A...AQ..
A. nancymaae            ..............................H...H..........I...G.TG.....Y........L..P.......N--------L.........RE.
S. boliviensis          ..............................H...R..........I.S.G.T.P....Y.....A..V..P....A..G--------LS........R..
R. norvegicus           EK...S...L.E........K............VH..VKSAGA..LP.P.TT.....A..L..PA..IK.PLW...V.N--------H....A-------
M. musculus             EK...P.....E........K............AR..IKSAGAR.HSDP.TT.R.I..Q.L.WPAT.T---VW.....N--------Q.....-------
H. glaber               ...............M...............P..A.SA...RV.SP..P..VI.S..G...T.VA.....P.WS.EF.N--------LS.R.I.......
F. damarensis           .K.M....I......I...............P..F.SA...RV..P..A..-..S..G...M.V....HIP.WS.EF.N--------L..H.I.......
O. degus                .Q......I.S..H.I...............P..F.SA...NV.DP..H...G.S.TG...V......HIQNWGREF.N--------L.RP.I..F..A.
C. lanigera             .Q......I.T....M...............P....ST...RL..L.SC.....S..G...P......HGP.W.SEFHN--------L.VP.........
C. porcellus            .K.....ST.HM...I...............P..N.SA...RV..P..P..NYS...S...T..S...H.PCWSSES.H--------V.R..A......I
I. tridecemlineatus     .K..................K.........H.N....V...R...L..PN....SS..N..T..N....GP.W.TD..T--------VP...V.....V.
M. marmota              .K..................K.........H.N....V...S...L..PN....SS..N..T.......GP.W.TD..N--------VP...I.....V.
O. cuniculus cuniculus  .K..R.P...S...S..........Y......T....V.K.GI.VHS.PGTT..S..A...T.......LP.W...V..--------L........E...
O. cuniculus algirus    .K..R.P...S...S..........Y......T....V.K.GI.VHS.PGTT..S..A...T.......LP.W...V..--------L........E...
S. floridanus           .K.MR.P...S...S..........Y......T....V.K.GI.VHS..GTT.PS..A...T.......PP.W...V..--------L........E...
S. bachmanni            .K.MR.P...S...S..........Y....P.T....V.K.GI.VHS.PGTT..S..A...T.......PP.W...V..--------L........E...
L. americanus           .K.MR.P...S...S..........Y....P.T....V.K.GI.VHS.PGTT..S..A...T.......PP.W...V..--------L........E...
L. timidus              .K.MR.P...S...S..........Y......T....V.K.GI.VHS.PGTT..S..A...T.......PP.W...V..--------L........E...
L. europaeus            .K.MR.P...S...S..........Y......T....V.K.GI.VHS.PGTT..S..A...T.......AP.W...V..--------L........E...
L. granatensis          .K.MR.P...S...S..........Y......T....V.K.GI.VHS.PGTT..S..A...T.......PP.W...V..--------L........E...
O. princeps             .K.MR.....S...S.................T.H..I..PGLQVPS.LG.VNQCSVL...VR.N..SPPP.W...F..--------L..L........A
O. collaris             .K.MR.....S...S.................T.H..I..PGLQVPS.LG.VNQCSVL...VR.N..SPPP.W...F..--------L..L........A
M. brandtii             .K........G....................P....TV..PGM..L.S.NT..P.......A...C.S.G.HWT--P.N--------PYD.RA...GA..
```

```
E. fuscus               .K........G....................P....AV..PGV..L.S.NT..P.S.....A...C.SPG.HWT--P.N--------PYD.RA...GA..
M. natalensis           ..........MV...................P..H.TV.IPGV..L.A.N...PR.T...AV..A....APPW...P.N--------PYDLR.GS.GA..
D. rotundus             ......P........................P...Q---.LGV.DL.S.N.T....E....V.....T.G.HW...S..--------P...R..S.GL..
H. armiger_             ..............S................P....V..PGI..L.ST..TT........T..........W..A.FG--------T...R.....LQ.
P. alecto               .G.............................PK...V..PRI..L.S...T.........T........V.W...S..--------P.........L..
R. aegyptiacus          .G....S.K.M..D.................PK...V..PGI..L.S...T.........T.........NW...S..--------P.........L..
M. putorius             .....-PL....I..................PK...V..PGV-.LP....T.P...L...T.P........W....NN--------PFS.....Q.L.A
E. lutris               .....-PL....V..................PK...V..PGV-.LT....T.P...L...T.P........W.....N--------PFP.....Q.L.A
C. ursinus              .....-PLK..........................HV..PGV..LP....T.P...L...T..........W.....N--------PFA.....Q.L..
Z. californianus        .....-PLK..........................HV..PGV..LP....T.P...L...T..........W.....N--------PFA.....Q.L..
O. rosmarus             .....-PLK..........................HV..PGV..LP....T.P...L...T..........W.....N--------PFA..S..Q.L..
A. melanoleuca          .....-PL..........................H..V..PGV..LP....T.P.S.L..ST.........W.....N--------PFT.....Q.L..
U. maritimus            .....-PL..........................H..V..PGV..LP......P...L..ST.........W.....N--------PFT.....Q.L..
C. lupus                .....-PL..S..........................I..PGV..LP..I.T.P......K......P.R.W..E...--------PFN.R....AL..
V. vulpes               .....-PL..S..........................I..PGV..LP..I.T.P......T......P.R.W..E...--------PFK.R....AL..
A. jubatus              .....-PL...MV...................K...V..PGV..IP....TG.......T..........W.....N--------LY.L.A....L..
F. catus                .....-PL...MV...................K...V..PGV..IP....TG.......T..........W.....N--------LY.L.A....L..
P. pardus               .....-PL...M....................K...V..PGV..IP....TG.......T..........W.....N--------LY.L.A....L..
E. asinus               ...................................D.V..SGI..LT.P..N..A.....T.....H.PHWH..S..--------P.........L..
E. caballus             ...................................D.V..SGI..LTHP..N..A.....T.....H.S.GH..S..--------P.........L..
C. simum                .K.........M.......................D.V..SGI..L...G.N.Y......A.....H.P.W...S..--------P..L.....L..
B. taurus               ..R..E...P.M..SS......................A.SPGI..L..PGVM..TS.AQ..A..S....SP.WRD.SF----------.PL.V....L..
B. bubalis              ..R..E...P.M..SS....K.................A.SPGI..L..PGVM..TS.AQ..A..S....SP.WR..SF----------.PL.V....L..
P. hodgsonii            ..R..E...P.MT.SS...G..................A.SPGV..FT.PGVM..TS.AQ..A..S....SP.WRDVSFP--------LA.L.V....L..
C. hircus               ..R..E...P.MTHSS......................A.SPGI..F..PGVI..TS.AQ..A..S....SP.WRDASFP--------LP.L.V....L..
S. scrofa               ..R..E..GGSA..GG.................H.PA.SPAV..L.CAG---P.S.AQH.A..S.H..P...SH.SHK--------PQ.L.A....L.A
C. dromedarius          ..R..E.....A..S....K.................AA..PGI..P.C.G.T.....T...A......RL..WS..SC.--------P.........L.A
C. ferus                ..R..E.....A..S....K.................AA..PGI..P.C.G.T.....T...A..I...RL..WS..SC.--------P.........L.A
V. pacos                ..R..E........S....K.................AA..PGI..PTC.G.T..S..T...A......HL..WS..SC.--------P.........L.A
O. orca                 ..H.EE.......DR.Q.....................A..AGL..L...G..N..S..Q..A.......LPHWR...FT--------.L...A..V.LR.
L. obliquidens          ..H.EE.......DR.Q.....................A..PGL..L...G.TN..S..Q..A.......LPHWR...CT--------.L...A..V.LR.
T. truncatus            ..H.EE.......DR.Q.....................A..PGL..L...G.TN..S..Q..A..A....LPHWR...CT--------.L...A..V.LR.
G. melas                ..H.EE.......DR.Q.....................A..PGL..L...G.TN..SV.Q..A.......LPHWR...CT--------.L...A..V.LR.
N. asiaeorientalis      ..H.EE.......DR.Q...............G.....A..PGL..L...G.T...S..Q..A.......L..WR...CT--------.L...A..V.LR.
P. sinus                ..H.EE.......DR.Q...............G.....A..PGL..L...G.T...S..Q..A.......L..WR...CT--------.L...A..V.LR.
M. monoceros            ..H.EE.......DR.Q.....................A..PGL..L...G.TN..S..Q..A.......L.HWR...C.--------.L...A..V.LR.
D. leucas               ..H.EE.......DR.Q.....................A..PGL..L...G.TN..S..Q..A.......L.HWR...CT--------.L...A..V.LR.
L. vexillifer           ..R.EE.......DR.Q.....................A.SPGL..L...G..N.....Q..T.......L..YR...CT--------.L...A..V.LRN
P. catodon              ..H.EER.GWD..DSGQ..................C.DARRPGL..L.NPG..H..G..Q..A.......L..WR...CP--------.P.L.A..V.LRA
B. musculus             .GH.GG..GR...DS.Q.................S..A..PGL..L...G.......Q..T.......L.SWR...CT--------PL...AS.V.LR.
B. acutorostrata        .GH.GR..GR...DS.Q.....................A..PGL..L...G.......Q..T.......L.SWR...CT--------PL...AS.V.LR.
L. africana             ..KE.W.............K.................T..PGV..I..P..T..VDVFQ.LP.......LP.W.S....SS------P..S.A.T....I
T. manatus              ..KE.W.............K.................T..PGV..L..PT.T..TDVFP.L..........P.W.....NS-------P..S.A......I
E. edwardii             ..KE.W----.........K..........P.L...A..PGM..L.S....N.SDR.LHTP.......IT.W.S..HNNQYNSYTTVNQ--I.TP..R.
E. telfairi             ----------------------------------------------------------------------------------------------------
C. asiatica             ..TV.G.......F.....K..........P.T...ELKPGV..P....TTC..D.FQ.QP.P.N....P.W..R..NH---------P.TLGTRLHDP

                                  510       520       530       540       550       560       570       580       590       600
                        ....|....|....|....|....|....|....|....|....|....|....|....|....|....|....|....|....|....|....|....|
H. sapiens              AGPRVWYRPIPSHMPSLHNIPVPETNYLGNTPTMPFSSLPP-TDESIKY--TIYNSTGIQIGAYNYMEIGGTSSSLLDSTNTNFKEEPAAKYQAIFDNTT
P. troglodytes          .........................................-.......--..................................S..........
P. paniscus             ....................................G....-.......--.............................................
P. anubis               .......G....P.S..YK.......V........T...L-........--A................V...........M......S..........
M. mulatta              .......G....P.S..YK................T...L-........--A................V..................S..........
M. fascicularis         .......G....P.S..YK................T...L-........--A................V..................S..........
R. roxellana            .D.....GLL..P.S...K.......V........T...L-....V...--A................V..................S..........
C. capucinus            .......G.....P...YKS.....F.............L-P.....C--..F..S...........V..M....S.N..V.......S...D.....
A. nancymaae            .......G.....L...YKS.....F............S.-P.....C--..F..S...........V..M....S....M.......S...D.....
S. boliviensis          .......G.....L..QYKS.....F..............-PE....C--..F..R...........V..M....S....V.......S...D.....
R. norvegicus           T.TG...G.SV.QSYNAYKT.....LP.SI....YI..A.-P.D..RC--......SRV..N....DL-.P..QPPTN.C---.V.STSRH..V.A...
M. musculus             T.TG...P.NL.Q.Y.TYKT.....IP.S.....YF.G.V-A.DL...--...F..S.....NH...DV-.LN.QPPNN.C---...STSRH.......
H. glaber               P.SGI..ALH..Q..N.FKT.....PA.S...L..Y...S-...LM.C--N.S.........T........MN.Q.DNV-C..LE..S.S.........
F. damarensis           S.--I..ALH..QI...FKT...SSGA.S...L..Y...S-...LM.C--N.S.........T........M..Q.-.N----LE..S.S.........
O. degus                P.AGTC.G.H..Q.A..FKT.....LA.S...L.LFCFSS-...L..SQVNVS..SN..F........RRMN.QP-EN.C..LE..STS.........
C. lanigera             S.AGI..G.HS.Q.SN.SKT.....LA.....L..Y.FIS-P..L..SHIN.S..S...F........TPVN.Q.-EN.R..LE..S.S......A...
C. porcellus            PST-..FGSHLNQ..NPFTP.M...SLA.S...L.LY.WM.-P..LLRSHIN.S......F.TS.....S.MNAP.-EN.C..VE..S.S.........
I. tridecemlineatus     .STGF..GLNS.P..N.YKT.....LPA....I.LY.F.S-....P.C--C..........T.....V.AVN.Q.PEN.CA.V....LS.FRD.....S
M. marmota              .STGF..GLNS...SN.YKT.....LPAS...I.LY.F.S-....P.C--C................V.AVN.Q.PENICA.V....LS.FRD.....S
O. cuniculus cuniculus  ..QA...G.SL.P.S..YKT.....L......I.....S.-AE..M.C--S....S...L..C.......I..PP.E-CLCTNLQ...S.......D..
O. cuniculus algirus    ..QA...G.GL.P.S..YKT.....L......I.....S.-AE..M.C--.....S...L..C.......M..PP.E-CLCTNLQ...S.......D..
S. floridanus           T.QPI..G.SL.P.S..YKT....S.L.....I.....L.-PE..V.C--.....S...L..C.......M..-P..-CSCTGLQ...S.......D..
S. bachmanni            T.QT...G.SL.P.S..YKT.....L......I.....S.-AE.AM.C--S....S...L..C.....S.V..-P..-CSCTGLQ...S.......D..
L. americanus           T.QT...G.SL.P.S..YKT.....L......I.....S.-AE.AM.C--S....S...L..C.......M...P.E-CSCTHLQ...S.......D..
L. timidus              T.QT...G.SL.P.S..YKT.....L......I.....S.-AE.AM.C--S....S...L..C.......M...P.E-CSCTHLQ...S.......D..
L. europaeus            ..QT...G.SL.P.S..YKT.....L......I.....S.-AE.AM.C--S....S...L..C.......M...P.E-CSCTHLQ...S.......D..
L. granatensis          ..QT...G.SL.P.S..YKT.....L......I.....S.-AE.AM.C--S....S...L..C.......M...P.E-CSCTHLQ...S.......D..
O. princeps             V..S...ASAG.QIQ.Q.KT.....L......I..T..A.-AE....C--.....S......C.......GL.P.S.-GLYANPQ..PL.DH....DA.
O. collaris             V..S...ASAG.QIQ.Q.KT.....L......I..T..A.-AE....C--.....S......C.......GL.P.S.-GLYANPQ..PL.DH....DA.
M. brandtii             IDS.F.SG.NA.QI....KT.....L......I..I..TS-R..TA..--A....A.V...DN.R..V-.V.LP.....YV.WN-...S...DT.....
E. fuscus               G.S.F.SG.DANQ..G..KT....TL......I..I..TS-R..TA.C--A....A.V...DN.R..V-.VNLP.....YV.W.-...S...Y......
M. natalensis           ....F..G.NA..T.G..KT.....L.E....I..I.RTS-RG.TAQ.--S....A.V...DS.H..V..M.LPFP.Y.YM.W.-...SA..D......
D. rotundus             V.S..CCG.N........KT.....L......I..V..SS-.G..V.C--S...........DK....V..V..P.....YI.L.-...S...D......
H. armiger_             V.S..CNG.N..Q.S..PKT.....LP.....I..V.F.S-R...VRC--S......V...DN.H....RMN.WS....YM.L.....S..LD..G...
P. alecto               V.S....G.T........KN.....L...L..I..I...S-R...V.C--S....A.....DN....V.ER..L.....Y..S...SIS..LD..G...
R. aegyptiacus          V.S....G.T........KS.....L...L..I..I..SS-G...V..--S....A.V...DN....V.....P.P.N.Y..S....IL..LD..G...
M. putorius             .SS.A..G.N.N....V.KT.....L....A.LS.G.F.S-R...V.C--S.R..A......DH.....S.M..VP..---M..........HD..E.I.
E. lutris               .SS.A..G.N......V.KT.....L......LS.ATF..-R...V.C--S.H.........EH.....S.M..VP..---M.L.........HD..E.I.
C. ursinus              T.S..L.G.N......V.K......L......L.C..F.S-R...V.C--.....A...V.DH.....S.M..V.P.---M.L.....S...D..E...
```

```
Z. californianus        T.S..L.G.N......V.K.......L......L.C..F.S-R...V.C--.....A...V.DH.....S.M..V.P.---M.L.....S...D..E...
O. rosmarus             T.S..L.G.N......V.K.......L......L....F.S-R...V.C--.....A...V.DH.....S.M..V.P.---M.L.....S...D..E.I.
A. melanoleuca          ..S....G.N......V.KT......L......I..T.F.S-R...V.C--A....AS..V.DH.......MN.V.P.---M.L.....S...D..E.V.
U. maritimus            ..S....G.N......V.KT......L......I..T.F.S-R...V.C--A....A...V.DH.......MN.V.P.---M.L.....S...D..E.V.
C. lupus                T.S....Q.N...I..VNKT......L......I....F.S-R...V.C--.....S...V.EH....V..M..A...---M.V.....Y...D....V.
V. vulpes               T.S....Q.N...I..VNKT......L......I....F.S-R.....C--.....S...V.EH....V..M..A...---M.V.....Y...D....V.
A. jubatus              PS.....G.N......V.KT......L......I......S-R.....C--N....A...V.DH.......LN.A..E..HV.L.....S...D....I.
F. catus                PS.....G.N......V.KT......L......I......S-R.....C--N....A...V.DH.......LN.A..E..HV.L.....S...D....V.
P. pardus               PS.....G.N......V.KT......L......I......S-R.....C--N....A...V.DH.......LN.A..E..HV.L.....S...D....I.
E. asinus               ....A..GAN....S.M.KT......L......I..I.F.S-R...A..--...........D........MT.Q...G.YM.L.Q...S...D....M.
E. caballus             ....AR.GAN....S.M.KT......L......I..I.F.S-R...A..--..C........H........MT.Q...G.YM.L.Q...S...D....M.
C. simum                ..Q....V.N....S.V.KT......V......V..I.F.S-R...A.C--...........DN.......M..Q.....YM.L.....S...D......
B. taurus               MS.....G.N.G.....YKT......L......I..T...S-R...S..--..HS.S.....DS.......M...V....YM.L.....S..........
B. bubalis              MS.....G.N.G.....YKT......L......I..T...S-R...S..--..HS.S.....DS.......M...V....YM.L.....S..........
P. hodgsonii            TS.....T.N.G.....YKT......L..S...I..T.V.S-R...A..--..H..S.....DS.......M...V...MYM.L.....S.......T..
C. hircus               TSS....A.N.G.....YKT......L......I..T.V.S-R...A..--..H..S.....DS.......M...V...MYM.L.....S.......T..
S. scrofa               .DR....M.SQG.R...YKT.....DL......V..A...S-R.A.PR.--IL.GNS.V...DH....L..A.V.V...MCPHV.....S......E...
C. dromedarius          .........N.G.....YKT......L......I......S-R...V..--..HS.S.....DH......A....V...MFV.V.....S...D..G.S.
C. ferus                .........N.G.....YKT......L......I......S-R...V..--..HS.S.....DH......A....V...MFV.V.....S...D..G.S.
V. pacos                .........N.G.....YKT......L......I......S-R...V..--..HG.S.V...DH......A....V...MFM.V.....S...D..G.SA
O. orca                 ...G.GDG.N.G.....YKT......L...........F.S-R...L..--G....F.V...NN.F.....M.PLV....YM.L..D..S..RD......
L. obliquidens          ...G..DG.N.G.....YKT......L......I....F.S-R...L..--G...CS.....NN.F.....M.PLV....YM.L..D..S..RD...D..
T. truncatus            ...G..DG.N.G.....YKT......L......I....F.S-R...L..--G...CS.V...NN.F.....M.PLV....YM.L..D..S..RD......
G. melas                ...G..DG.N.G.....YKT......L......I....F.S-R...L..--G...CS.V...NN.F.....M.PLV....YM.L..D..S..RD......
N. asiaeorientalis      ...G..DG.N.G.....YKT......L....S.I....F.S-R...L..--G..KCS.V...NN.......M.PLV....YM.L.GD..S...D......
P. sinus                ...G..DG.N.G.....YKT......L.E..S.I....S.S-R...L..--S...CS.....NN.......M.PLV....YM.L.GD..S...D......
M. monoceros            ...G..DG.N.G.....YKT......L......I....F.S-R...L..--G....F.V...NN.......M.PLV....YM.L..D..S...D......
D. leucas               ...G..DG.N.G.....YKT......L......I....F.S-R...L..--G....F.V...NN.......M.PLV....YM.L..D..S...D......
L. vexillifer           T..G..NG.N.G....MYKT......L......I......S-RGK-------------------------------------------------------
P. catodon              ...G.CCG.H.G.T....KT.....HP......I......S-R...L..--S....S.V...KN.......MG.LV..G.YMSL.DA..SQ..D......
B. musculus             ...G...G.N.G.....YKT......L......I......S-R...L..--S....S.....NN.......M..VV....HM.L..G..S...D......
B. acutorostrata        ...G...G.N.G.....YKT......L......I......S-R...L..--S.C..S.....NN.......M..VV....HM.L..G..S...D......
L. africana             ....P..G.N.....N..KT......L......I...-F.S-I...FR.--S..........N.....V.RM.VPP.......L.....W...E....I.
T. manatus              ....P..G.N........KT.....SL.V....I..I.F.S-....FR.--N......V...N.....V.RM.VPPQ............S...E....I.
E. edwardii             S...P.FE.N...I.P..KT.....SL.SFHLPTIPFQSFSS....TR.--N.H..S.....NN.F.....AHTQP.E..SV.Y-....L...E..A.N.
E. telfairi             ------------------------------------------------------------------------------------------------..LS
C. asiatica             ETGGPGHWCN.TQV....KS....S.L......I....Y.S-.EN.VR.--N....S.....D.SF..V.RL.VPP....Y........S...E..G.V.

                                610       620       630       640       650       660       670       680       690
                        ....|....|....|....|....|....|....|....|....|....|....|....|....|....|....|....|....|....|..
H. sapiens              SLTDKHLDPIRENLGKHWKNCARKLGFTQSQIDEIDHDYERDGLKEKVYQMLQKWVMREGIKGATVGKLAQALHQCSRIDLLSSLI-YVSQN
P. troglodytes          ......................................................................................-.....
P. paniscus             ......................................................................................-.....
P. anubis               .......................................................................V....A.....NH..-.....
M. mulatta              .......................................................................V....A.....NH..-.....
M. fascicularis         .......................................................................V....A.....NH..-.....
R. roxellana            ............................................................................A.....NY..-.....
C. capucinus            ................Q.........L............................L..........................NH.V-.I...
A. nancymaae            ................Q.......................................R.........................NH.V-.....
S. boliviensis          ................Q......................................L....V.............R.......NR.V-.....
R. norvegicus           .......N.....M..Q...........E..........................L....T...............C.T...NQ..-QA..S
M. musculus             ....E..N.......RQ...........E..........................L....T...............C.....NH..-RA..S
H. glaber               ....N..N.V......Q...........E...............H...H......L....S.........W..Y....K...NC..-HIT..
F. damarensis           ....S..T.V......Q...........E...............H...H......L....S.........S..Y....K...TG..-HT...
O. degus                ....NQ.NL.G.....Q..P........EP..............H...F......R....SQ........W..Y..F.K...NH.V-DRG.I
C. lanigera             ...ANQ.T.VE........P........E...............H...V......R....S.R.......W..Y....K..VNH.M-DI...
C. porcellus            .V..I..N.V......Q..T......I.E...............H....K.....E....N.........R..Y....K...NH..-DI.K.
I. tridecemlineatus     ...A...N.V..................E..........................L....N.....................TA..-H....
M. marmota              ...A...N.V..................E..........................L....N.....................NA..-H....
O. cuniculus cuniculus  .......S.V.D..............L.E..........................L....T.........R.......Q...TC..-QL..K
O. cuniculus algirus    .......S...D..............L.E..........................L....T.........R.......Q...TC..-QL..K
S. floridanus           .......S.V.D..............L.E..........................L....T.........R.......K...TC..-QL..K
S. bachmanni            .......S...D..............L.E..........................L....T.........H.......K...TC..-QL..K
L. americanus           .......S.V.D..............L.E..........................L....T.........R.......K...TC..-QL..K
L. timidus              .......S.V.D..............L.E..........................L....T.........R.......K...TC..-QL..K
L. europaeus            .......S.V.D..............L.E..........................L....T.........R.......K...TC..-QL..K
L. granatensis          .......S.V.D..............L.E..........................L....T.........R.......K...TC..-QL..K
O. princeps             ....T..S.V.D.......T........E..........................L....T.........R..Q.SA.....NY.V-RL..K
O. collaris             ....T..S.V.D.......T........E..........................L....T.........R..Q.SA.....NY.V-RL..K
M. brandtii             .........V.....RT...........E..........................L....S.........R..Y........N..V-FI.M.
E. fuscus               .........V.....RT...........E..........................L....S.........W..Y........NF.G-TI...
M. natalensis           ....E....V.....RT..S....M...E..........................L....S.R.......R..Y........NC.VAHI...
D. rotundus             .........V.....RS..S........E..........................L....N.........Y.......L...NA.K-.I..H
H. armiger_             .........V.D....S..S........E..........................L....N.........YS.Y.......VN...-HI...
P. alecto               .........V.....RS..I.......SE..........................L....N........TY..Y....T...NA.V-.I...
R. aegyptiacus          .........V.....RS..S......ISE..........................L....N.........Y..Y..A.T...NA.V-.I...
M. putorius             .........V......Q...........EP.........................Q....N.........Y...........NY.V-HI..S
E. lutris               .........V......Q...........EP.........................Q....N.........Y..Y....T...NY.V-RI..S
C. ursinus              .........V......Q..........SEP..E...................KT.L....N.........Y..Y........NY.V-HIN.S
Z. californianus        .........V......Q..........SEP..E...................KT.L....N.........Y..Y........NY.V-HIN.S
O. rosmarus             .........V......Q..........SEP..E...................KT.L....N.........Y..Y........NY.V-HIN.S
A. melanoleuca          .........V......Q...........EP.........D...............L....N.........Y..Y........NY.V-HI..S
U. maritimus            .........V......Q...........EP..........................L....N.........Y..Y........NY.V-HI..S
C. lupus                .........V......Q...........EP..........................L....N.........Y..Y........NY..-HI...
V. vulpes               .........V......Q...........EP..........................L....N.........Y..Y........NY..-HI...
A. jubatus              .........V.....RQ...........E..........................L....S.........Y..Y........NY..-HI..S
F. catus                .........V.....RQ...........E..........................L....S.........Y..Y........NY..-HI..S
P. pardus               .........V.....RQ...........E..........................L....S.........Y..Y........NY..-HI..S
```

| | |
|---|---|
| E. asinus | .........V...............S..E........................R.L....N.........Y...........NH..-HI..Y |
| E. caballus | .........V...............S..E........................R.L....N.........Y...........NH..-HI..Y |
| C. simum | .........V...............N..E..........................L....N.........C..Y........HR..-HI... |
| B. taurus | .........V.....R...........SE..........................L..A.S.........C..YL......VNC..-RI..S |
| B. bubalis | .........V.....R...........SE..........................L..A.S.........Y..YL......VNC..-RI..S |
| P. hodgsonii | .........V.....R...........SE..........................L....S.........F..YL......VNC..-RI... |
| C. hircus | .........V.....R...........SE..........................L....S.........L..YL......VNC..-R.... |
| S. scrofa | .........V......N..I......L.E..........................L....S.........R..Y....L....C..-RS... |
| C. dromedarius | .........V...............Y..E..........................L....N.........Y..Y.......VNY..-NI... |
| C. ferus | .........V...............Y..E..........................L....N.........Y..Y.......VNY..-NI... |
| V. pacos | .........V...............Y..E..........................L....N.........Y..Y.......VNH..-NI... |
| O. orca | .........V.....RQ..D.......S...........................L....S.........F..Y....L....H.T-HS... |
| L. obliquidens | .........V.....RQ..D.......S...........................L....S.........F..Y....L...RH.T-HS... |
| T. truncatus | .........V.....RQ..D.......S...........................L....S.........F..Y....L....H.T-HS... |
| G. melas | .........V.....RQ..D.......S...........................L....S.........F..Y....L...NH.T-HS... |
| N. asiaeorientalis | .........V.....RQ..D.......Y...........................L....S.........W..Y....V....H..-HS... |
| P. sinus | .........V.....RQ..D.......Y...........................L....S.........W..Y....V....H.T-HS... |
| M. monoceros | .........V.....RQ..D.......S...........................L....S.........W..Y....V....H.T-HS... |
| D. leucas | .........V.....RQ..D.......S...........................L....S.........W..Y....V....H.T-HS... |
| L. vexillifer | -------------------------------------------------------------------------------------------- |
| P. catodon | .........V..........F......S...........................L....S.........C...R...T...NC..-HA..D |
| B. musculus | ....R....V..........F......S...........................L....S......R..Y..YE...V...NC..-CM... |
| B. acutorostrata | ....R....V..........F......S...........................L....S.........C..Y....HSDT.--------- |
| L. africana | .........V.....R...K........EP.........................L....S.........Y..Y....T...NY..-R.... |
| T. manatus | .........V.....R...K........EP.........................L....N.........Y..Y....T...NY.V-R.... |
| E. edwardii | .........V......Q..K........EP..............R..........L.K..NR.......TY..YH......VNY.K-D.... |
| E. telfairi | .........V.........K........EP.........................L....N.........H..Y......I.HY..-VA... |
| C. asiatica | .........V.........K........EP.........................L....S.........Y..Y.......VN...-V...H |
