## Supplementary material for "Convergent loss of the necroptosis pathway in disparate mammalian lineages shapes virus countermeasures": S appendix: S Appendix 5.pdf

**S Appendix 5.** RIPK3 and MLKL protein alignment from species from rodent and afrotheria lineages.

**S Appendix 5.1. Rodent RIPK3 protein alignment.** RIPK3 protein alignment from human and 5 rodent genomes. Stop codons are indicated by an asterisk (*) and grey boxes.

```
                       10        20        30        40        50        60        70        80        90       100
               ....|....|....|....|....|....|....|....|....|....|....|....|....|....|....|....|....|....|....|....|
H. sapiens     MSCVKLWPSGAPA-PLVSIEELENQELVGKGGFGTVFRAQHRKWGYDVAVKIVNSKAISREVKAMASLDNEFVLRLEGVIEKVNWDQDPKPALVTKFMEN
M. musculus    ..S.....T..S.V....R...KKL.F.......V....H..T.NH..........K..W.....VN.R..N..L.L..T.DLQ..FVSGQ....R....
R. norvegicus  ..S....LN..SSIS..GS.....LGF.......A....R.TA.NL.........K.......VN.RH.N..L.L..T.NLE..YVYG.....G....
O. degus       .NRR-.ISVS.-SLVSSENLKNPDFVSE.GF.TVLRAQYRKWDCDAAAKIMHSDAISKEVKTITSLNN--QY....L.ETKSLQ.XGMSR.....QL...
F. damarensis  ----------------------------------------------------------------..NPHRQY....L..A.NLQ.EGVSG.....P.L..
H. glaber      ----------------------------------------------------------------..N.R.LY....L..T.NLQ.NGVSG.....Q....

                      110       120       130       140       150       160       170       180       190       200
               ....|....|....|....|....|....|....|....|....|....|....|....|....|....|....|....|....|....|....|....|
H. sapiens     GSLSGLLQSQCPRPWPLLCRLLKEVVLGMFYLHDQNPVLLHRDLKPSNVLLDPELHVKLADFGLSTFQGGSQSGTG----SGEPGGTLGYLAPELFVNVN
M. musculus    ...A....PE...........Q......C...SLD.P..........I.......A..................S.SGSG.RDS....A..D...LFK..
R. norvegicus  ........PS...........E......C...SL..S...............L..A..................S.SG--.RDS....A......LD.DG
O. degus       .......CRDW.G..LF..C..Q......N...SL.S....*....S....TN..A.V.........P......CE--TQQ.RD..A......LADI.
F. damarensis  S.......RD..X....V....L.M....S...IL.S...R...........TK..A............K......CR--FQ......A......LADIF
H. glaber      ........RD.......V....Q......S...NL....................TN..A........M.....L..A.YG--.QG..S..A......LADIF

                      210       220       230       240       250       260       270       280       290       300
               ....|....|....|....|....|....|....|....|....|....|....|....|....|....|....|....|....|....|....|....|
H. sapiens     RKASTASDVYS--FGILMWAVLAGREVELPTEPSLVYEAVCNRQNRPSLAELPQAGPETPGLEGLKELMQLCWSSEPKDRPSF-QECLPKTDEVFQMVEN
M. musculus    L...K......--....V.........A..VDKT..IR.T..D..S..P.T...PGS.......K.....IH..G.QSEN....-.D.E...N..YNL.KD
R. norvegicus  -...K......--..V.V.T......A.VVDKT..IRG......R..P.T...PDS...............TH............-.D.ES..NN.YIL.QD
O. degus       K...K......KI....I..L*....P.V-----------TKK.IQALMT...LSS.......E..K...RS..H..G..APL-PRMPGNNQKGLLGAHG
F. damarensis  K..TK......--....L*..X....P.VVPQI..IR.T.AEKRIQ.P.T...PSS.......E.EN...RF..H..S..LP.FPK.QET.KKA.*R.HD
H. glaber      K..TK......--.Q..........T.VVPQT..IH.T.VEK.I..P.T...PCR.......K.EK.I.C...H..N...R.FPGE.SFHWQLE*----

                      310       320       330       340       350       360       370       380       390       400
               ....|....|....|....|....|....|....|....|....|....|....|....|....|....|....|....|....|....|....|....|
H. sapiens     NMNAAVSTVKDFLSQLRSSNRRFSIPESGQGGTEMDGFRRTIENQHSRNDVMVSEWLNKLNLEEPPSSVPKKCPSLTKRSRAQEEQVPQAWTAGTSSDSM
M. musculus    KVD....E..HY...H...G.NL.AR.PS.R.....CP.E----------T...KM.DR.H....SGP..G..----PERQ..DTS.GP.TP.R....PV
R. norvegicus  KVD....K..HY...Y...DTKL.AR..S.K...V.CP.E----------TI.Y.M.DR.H....SG...ERLT...E.RGK-.ASFGH.TP......TL
O. degus       .VD..ILEHRICEP-.STVAAMAAKIV.QHKTPDISCYFS*------------------------------------------------------------
F. damarensis  K.SD.I.K.TK...EH......L.A..PS.R......GTT--GS.CTKIGS.L..S..C.R.KQF.T...ET.TGF.EER...K...QH.R......Y..
H. glaber      ----------------------------------------------------------------------------------------------------

                      410       420       430       440       450       460       470       480       490       500
               ....|....|....|....|....|....|....|....|....|....|....|....|....|....|....|....|....|....|....|....|
H. sapiens     AQPPQTPETSTFRNQMPSPTSTGTPSPGPRGNQGAERQGMNWSCRTPEPNPVTGRPLVNIYNCSGVQVGDNNYLTMQQTT-ALP--TWGLAPSGKGRGLQ
M. musculus    .GT..I.H.LP..GTT.G.VF.E..G.H.QR...DG.H.TP.YPW..-...M..P.ALVFN...E..I.NY.S.VAPPR.T.SSSAKYDQ.QF.R...W.
R. norvegicus  .GT..I.H.LPS.GTT.R.AF.E..G.D.QR...DG.NSNP.YTWNA-...M..LQSIVLN...E..I.QH.CMSV.PR.-.F.--KKEP.QF.R...W-
O. degus       ----------------------------------------------------------------------------------------------------
F. damarensis  ..TL.....LP.Q..TGR.IV.......AQ...VRNWRNCSLTRKNLGLRHRAHTLTQPVAGEFPLVFPPGEHEAPLSSSPVGLQTSSFRL*--------
H. glaber      ----------------------------------------------------------------------------------------------------

                      510       520
               ....|....|....|....|....|....
H. sapiens     HPPPVGSQEGPKDPEAWSRPQGWYNHSGK
M. musculus    PFHK-------------------------
R. norvegicus  -----------------------------
O. degus       -----------------------------
F. damarensis  -----------------------------
H. glaber      -----------------------------
```

**S Appendix 5.2. Afrotheria RIPK3 protein alignment.** RIPK3 protein alignment from human and 2 afrotheria genomes. Stop codons are indicated by an asterisk (*) and grey boxes.

```
                    10        20        30        40        50        60        70        80        90       100
            ....|....|....|....|....|....|....|....|....|....|....|....|....|....|....|....|....|....|....|....|
H. sapiens  ---------------------------------------------------------------------MSCVKLWPSGAPAPLVSIEELENQELVGKGG
L. africana ----------------------------------------------------------------------------------------------------
T. manatus  MQGGTVEAPVTCRRRKLLSPSVLRSDPGSETVPRLEIAPLQRLPPTPPPRSHPIRVSHDFQKPLTPSSLT.SS...S..GS....PNK....LKF..Q..

                   110       120       130       140       150       160       170       180       190       200
            ....|....|....|....|....|....|....|....|....|....|....|....|....|....|....|....|....|....|....|....|
H. sapiens  FGTVFRAQHRKWGYDVAVKIVNSKAISREVKAMASLDNEFVLRLEGVIEKVNWDQDPKPALVTKFMENGSLSGLLQSQCPRPWP--LLCRLLKEVVLGMF
L. africana -----------------MAGWWTWHL.QAQQLLRTPGAQRE.DSPSSSA.TGVGECVGS....*....T...E...H.R.G..L*L.R.L.PR...X---
T. manatus  ..A...H..A........LL.............N.R.QY..L.L..T..LE.ENVCG.T...R.........E...H......Q*Q.G....H......C

                   210       220       230       240       250       260       270       280       290       300
            ....|....|....|....|....|....|....|....|....|....|....|....|....|....|....|....|....|....|....|....|
H. sapiens  YLHDQNPVLLHRDLKPSNVLLDPELHVKLADFGLSTFQGGSQSGTGSGEP-GGTLGYLAPELFVNVNRKASTASDVYSFGILM-----WAVLA----GRE
L. africana ...NL......*G.........L..RA.....S.....*..FFFVR.RVR*V..A.S.......LAD..QR..MDTSRP.PSC.STQTILLPSH.PLDHTQL
T. manatus  ...SL......Q..........L...A............R..A.FS..-....S.......LAD..QR.TMD..........-----.S...----..

                   310       320       330       340       350       360       370       380       390       400
            ....|....|....|....|....|....|....|....|....|....|....|....|....|....|....|....|....|....|....|....|
H. sapiens  --------------------VELPTEPSLVYEAVCNRQNRPSLAELPQAGPETPGLEGLKELMQLCWSSEPKDRPSFQECLPKTDEVFQMVENNMNAAV
L. africana PFSMMKKGAYPQCLLFSSTPHPAVVIQTW.AQV.M.EK....L.S...EPR.N.L......K...H...HK........D.*S..S...LLEQDKRD...
T. manatus  --------------------A.MVVQT..AQV...EK.E..X.S...EL..N...X---.K...H...H..R.....HD.*S..S.A.LL.QDK.D...

                   410       420       430       440       450       460       470       480       490       500
            ....|....|....|....|....|....|....|....|....|....|....|....|....|....|....|....|....|....|....|....|
```

```
H. sapiens    STVKDFLSQLRSSNRRFSIPESGQGGTEMDGFRRTIENQHSRNDVMVSEWLNKLNLEEPPSSVPKKCPSLTKRSRAQEEQVPQAWTAGTSSDSMAQPPQT
L. africana   .M..K...EH.R....LFSL.P.R.E....DPGGIMGSLC.W..ST...S..S.H...C..T.L...T...E-I.T.R...QDTKI..A....TT*L...
T. manatus    .M..K...EQ.G....L.LL.P.P.ER...DPG.IMGS.C.W..SI...S..N.H...C.GT..E.ST...EKI.T.GG..QDTRI..A....T......

                      510       520       530       540       550       560       570       580       590       600
              ....|....|....|....|....|....|....|....|....|....|....|....|....|....|....|....|....|....|....|....|
H. sapiens    PETSTFRNQMPSPTSTGTPSPGPRGNQGAER--------QGMNWSCRTPEPNPVTGRPLVNIYNCSGVQVGDNNYLTMQQTTALPTWGLAPSGKGRGLQH
L. africana   .K..P..S.I.N.PQV.SQVL..KEIRDPI--LAFPQGEREVPLLRLILLLISLK.QLSIVLDG.Q...I.N....NILGRPT...Q.P..PSV...W.N
T. manatus    .KI.P..S.T.NS..VWV.D..TQ.....--------RHDK..PHWDS.L..IPAVYSPTWVARGADWKQQLHEH.RETHP.HGGPSTSQR.-------

                      610       620       630
              ....|....|....|....|....|....|..
H. sapiens    PPPVGSQEGPKDPEAWSRPQGWYNHSGK---*
L. africana   L.G.S.E...EE......S..E.KNVNCCTF*
T. manatus    -------------------------------*
```

**S Appendix 5.3. Rodent MLKL protein alignment.** RIPK3 protein alignment from human and 3 rodent genomes. Stop codons are indicated by an asterisk (*) and grey boxes.

```
                      10        20        30        40        50        60        70        80        90       100
              ....|....|....|....|....|....|....|....|....|....|....|....|....|....|....|....|....|....|....|....|
H. sapiens    ----------------MENLKHIITLGQVIHKRCEEMKYCKKQCRRLGHRVLGLIKPLEMLQDQGKRSVPSEKLTTAMNRFKAALEEANGEIEKFSNRSN
H. glaber     MPEIVDFNKFHPLLPG.DK.GQ..S...Q.L.QW..F...QN.SQ..RDH.S..LQV.QW.....T.NLSP.ITAVLD.FQN-...K.MEKM...NTQTI
O. degus      ----------------------------------------------------------------------------------------------------
F. damarensis ----------------------------------------------------------------------------------------------------

                     110       120       130       140       150       160       170       180       190       200
              ....|....|....|....|....|....|....|....|....|....|....|....|....|....|....|....|....|....|....|....|
H. sapiens    ICRFLTASQDKILFKDVNRKLSDVWKELSLLLQVEQRMPVSPISQGASWAQEDQQDADEDRR---AFQM--LRRDNEKIEASLRRLEINMKEIKE-TLRQ
H. glaber     FRKV.MPGSN.....E..QM.R...EVFM.Q..ID.HVCI.S..K.EF.P....S..EK...FLL...SLKEVKSCLPRAR.ACLHARFSAWTLAWRRCC
O. degus      ----------------------------------------------------------------------------------------MKATMRWSEVDF
F. damarensis ------------------------------------------------------------MIHAYNPRTPVAEDGGLPQHTNTDV.HFSTSDSEGNHYNK

                     210       220       230       240       250       260       270       280       290       300
              ....|....|....|....|....|....|....|....|....|....|....|....|....|....|....|....|....|....|....|....|
H. sapiens    YL------PP-KC-MQEIPQEQIKEIKKEQLSGSPWILLRENEVSTLYKGEYHRAPVAIKVFKKLQAGSIAIVRQTFNKEIKTMKKFESPNILRIFGICI
H. glaber     RAAATVVLA.LGFVRR.P.GARDAPAPPHGRALRKLPVHVRLVLTEDERSLPDMTSLVVWAMAVDISYISVYDH.ARFTPHQIVR.HFHKE.SHMRKFDS
O. degus      KEIMETMKQYSLRPAYQTA.AK...M.E.E.L.FS*T.I.QSKF.KP......P.......NQ...KR.RTA.EH.H.-SSAV..SD..KN.H.....T
F. damarensis NMEATLRWLEVDLK*IKETLSRTNK..E.E.L.F..T...QS.F.K....A..X--....I.N..*TKC.GK.KEH.XT.TSAL..SH...N.H.CEV..

                     310       320       330       340       350       360       370       380       390       400
              ....|....|....|....|....|....|....|....|....|....|....|....|....|....|....|....|....|....|....|....|
H. sapiens    DETVTPPQFSIVMEYCELGTLRELLDREKDLTLGKRMVLVLGAARGLYRL-------------HHSEAPELHGKIRSSNFLVTQGYQVKLAGFELRKTQT
H. glaber     PNILXTFGIC..ERGSSPQFCMVMEYC.CRSLRDVLDEDRNLQL-..-.ILLALGAAKGFYWL...GG.H.QRN.S..S....KS.GSE---GFELREAQ
O. degus      VL.GSS....M.A.F.KHS....VWVK....S..LCFL.....G-NXL*GL------------YC.R.-S..RN.SC.S....R..K..-LIEFEL.QT*
F. damarensis V..GSL....M.T....CS..K.V..K.R..S..LCVL.L....K.....-------------..CK.LP..RNSS..S....A..E...-TGFEMRETQ

                     410       420       430       440       450       460       470       480       490       500
              ....|....|....|....|....|....|....|....|....|....|....|....|....|....|....|....|....|....|....|....|
H. sapiens    SMSLGTTREKTDRVKSTAYLSPQELEDVFYQYDVKSEIYSFGIVLWEIATGDIPFQGCNSEKIRKLVAVKRQQEPLGEDCPSELREIIDECRAHDPSVRP
H. glaber     TSISQKIK.TK-----...I...R.NNP.HK..I.A..............R...E....KE.CQ..CE.QL.QMRSK..SPL.Q.V....Q.YE..AW.
O. degus      TFISQK.KQNKTXQTTAYFSPQKLE-NLLRE*.IIA.....R...*.T...K...ES.A..EF*...FENP..GSKTQTHRVKAATLQL-----------
F. damarensis TSISQK.KGIEQSNPTEYFSPQNLE-NLLCEK.TIAK.............K..*GWAFK.ICQLVFKNPQ.DLLREDYP.QL*EVTDEFRASEPSGQPS

                     510       520       530
              ....|....|....|....|....|....|....|..
H. sapiens    SVDEILKKLSTFSK----------------------*
H. glaber     .M.SKALPTERGMSPVFLSQL---------------*
O. degus      ------------------------------------*
F. damarensis GDEILE.TVDFCCVACKNLKQRVLDKRLKETQTGHL*
```

**S Appendix 5.4. Afrotheria MLKL protein alignment.** MLKL protein alignment from human and 2 afrotheria genomes. Stop codons are indicated by an asterisk (*) and grey boxes.

```
                      10        20        30        40        50        60        70        80        90       100
              ....|....|....|....|....|....|....|....|....|....|....|....|....|....|....|....|....|....|....|....|
H. sapiens    MENLKHIITLGQVIHKRCEEMKYCKKQCRRLGHRVLGLIKPLEMLQDQGKRSVPSEKLTTAMNRFKAALEEANGEIEKFSNRSNICRFLTASQDKILFKD
T. manatus    .DE..Q..S...LVYQQ......WN..Q...KHIH..LQ........QKNL-.TQ..A.LLS.QTV....KDQ.K..N.K..VQK....GT.....SA
C. asiatica   .DT..Q......LVYNQ.....C.RH..Q...N.IQH.L...Q......EKNL-.VQ..D.LHH.QTI....KMR......K...LK..K.RD.....SA

                     110       120       130       140       150       160       170       180       190       200
              ....|....|....|....|....|....|....|....|....|....|....|....|....|....|....|....|....|....|....|....|
H. sapiens    VNRKLSDVWKELSLLLQVEQRMPVSPISQGASWAQEDQQDADEDRRAFQMLRRDNEKIEASLRRLEINMKEIKETLRQYLPPKCMQEIPQEQIKEIKKEQ
T. manatus    ..KR.R..SE....V...D..T------FHQP.QK...R..E..MIFAALFPAEK.N.DLL..*..S...-.I.......-K.PINKL........E...
C. asiatica   L.KR.E..SQ..L.V..AD..KLILNTLHRG..QE......K..MG..*RVE.QMGENIEF.L.QLEKN---I.......-Q.P.KQL...........

                     210       220       230       240       250       260       270       280       290       300
              ....|....|....|....|....|....|....|....|....|....|....|....|....|....|....|....|....|....|....|....|
H. sapiens    LSGSPWILLRENEVSTLYKGEYHRAPVAIKVFKKLQAGSIAIVRQTFNKEIKTMKKFESPNILRIFGICIDETVTPPQFSIVMEYCELGTLRELLDREKD
T. manatus    I.....Q..K..D.........KC..T..A.SNP..T..GT.....N..R.....D...V..........KEN.A..C.I..H..F........K.QN
C. asiatica   .L.A...E..K..Y........SKC..T....NNS.-S..GL..S..KN..R.....D...............EK.TK.C.I..H.D.........K.QN

                     310       320       330       340       350       360       370       380       390       400
              ....|....|....|....|....|....|....|....|....|....|....|....|....|....|....|....|....|....|....|....|
```

```
H. sapiens   LTLGKRMVLVLGAARGLYRLHHSEAPELHGKIRSSNFLVTQGYQVKLAGFELRKTQTSMSLGTTREKTDRVKSTAYLSPQELEDVFYQYDVKSEIYSFGI
T. manatus   .EF.VCIF.SX.............E....RN.S.TS...AE..H........S.....I.RKVKERRAEK.N....V...G.KN.YHK..I.A.......
C. asiatica  .EF.V.IF.A.............AE....RN.S.T.....G..H........T.....I.GQVKEKRAE..N....F...V.KN..HK....A.......

                     410       420       430       440       450       460       470       480
             ....|....|....|....|....|....|....|....|....|....|....|....|....|....|....|....|....|
H. sapiens   VLWEIATGDIPFQGCNSEKIRKLVAVKRQQEPLGEDCPSELREIIDECRAHDPSVRPSVDEILKKLSTFSK-------------*
T. manatus   ........K.L.K..D.RR..E.AESDGY.........P..Q....G...YE..E..L..VRAFSGRSIFQLQRARATHVPSHL*
C. asiatica  ........N......G.R..QE..KSGGCP....D......Q.V..G.Q.YE.AE.....GYPA--------------------*
```
