## Supplementary material for "Convergent loss of the necroptosis pathway in disparate mammalian lineages shapes virus countermeasures": S appendix: S Appendix 6.pdf

### **S Appendix 6.** Tables identifying RIPK3 and MLKL mutations and premature stop codons in Cetacea order.

### **S Appendix 6.1.** Table identifying RIPK3 mutations and premature stop codons in Cetacea order.

| | **Exon 1** | **Exon 2** | **Exon 3** | **Exon 4** | **Exon 5** | **Exon 6** | **Exon 7** | **Exon 8** | **Exon 9** | **Exon 10** |
|---|---|---|---|---|---|---|---|---|---|---|
| *Tursiops truncatus* | 7nt del (242); Stop (411) | Exon Not Found | Exon Not Found | Exon Not Found | Exon Not Found | OK | 1nt del (20); Stop (42) | OK | OK | 1nt del (12) |
| *Globicephala melas* | 7nt Del (242); Stop (411) | Exon Not Found | Exon Not Found | Exon Not Found | Exon Not Found | OK | 1nt del (20); Stop (42) | OK | OK | Exon Not Found |
| *Lagenorhynchus obliquidens* | 7nt Del (242); Stop (411) | Exon Not Found | Exon Not Found | Exon Not Found | Exon Not Found | OK | 1nt del (20); Stop (42) | OK | OK | Exon Not Found |
| *Orcinus orca* | 7nt Del (242); Stop (411) | Exon Not Found | Exon Not Found | Exon Not Found | Exon Not Found | OK | 1nt del (20); Stop (42) | OK | OK | 1nt del (12) |
| *Neophocaena asiaeorientalis asiaeorientalis* | Stop (84); 7nt Del (242); Stop (411) | Exon Not Found | Exon Not Found | Exon Not Found | Exon Not Found | OK | Stop (42) | OK | OK | 1nt del (12) |
| *Phocoena sinus* | Stop (84); 7nt Del (242); Stop (411) | Exon Not Found | Exon Not Found | Exon Not Found | Exon Not Found | OK | Stop (42) | OK | OK | OK |
| *Monodon monoceros* | Stop (84); 7nt Del (242); Stop (411) | Exon Not Found | Exon Not Found | Exon Not Found | Exon Not Found | OK | Stop (42) | OK | OK | Exon Not Found |
| *Delphinapterus leucas* | Stop (84); 7nt Del (242); Stop (411) | Exon Not Found | Exon Not Found | Exon Not Found | Exon Not Found | OK | Stop (42) | OK | OK | Exon Not Found |
| *Lipotes vexillifer* | Stop (411) | Exon Not Found | Exon Not Found | Exon Not Found | Exon Not Found | OK | Stop (42) | OK | OK | Exon Not Found |
| *Physeter macrocephalus* | Stop (411); 1nt del (445) | Unknown (Fragmented Genomic Region, N's) | Stop (110) | OK | Ok | OK | Stop (42); 2nt del (71) | OK | OK | OK |
| *Balaenoptera acutorostrata scammoni* | Stop (228); Stop (411) | 2nt ins (5); | Stop (110) | OK | OK | OK | Stop (42) | OK | OK | OK |

### **S Appendix 6.3.** Table identifying MLKL mutations and premature stop codons in Cetacea order.

| | **Exon 1** | **Exon 2** | **Exon 3** | **Exon 4** | **Exon 5** | **Exon 6** | **Exon 7** | **Exon 8** | **Exon 9** |
|---|---|---|---|---|---|---|---|---|---|
| *Tursiops truncates* (**Delphinidae**) | Exon Not Found | 1nt ins (116) | 1nt del (138) | 1nt del (11) | OK | 1nt del (87); Stop (161) | 2nt del (53) | 1nt ins (213) | OK |
| *Globicephala melas* (**Delphinidae**) | Exon Not Found | 1nt ins (116) | 1nt del (138) | 1nt del (11) | OK | 1nt del (87); Stop (161) | 2nt del (53) | 1nt ins (213) | OK |
| *Lagenorhynchus obliquidens* (**Delphinidae**) | Exon Not Found | 1nt ins (116) | 1nt del (138) | OK | OK | 1nt del (87) | 2nt del (53) | 1nt ins (213) | OK |
| *Orcinus orca* (**Delphinidae**) | Exon Not Found | 1nt ins (116) | 1nt del (138) | OK | OK | 1nt del (87) | Stop (5); 2nt del (53) | 1nt ins (213) | OK |
| *Neophocaena asiaeorientalis asiaeorientalis* (**Phocoenidae**) | Exon Not Found | 1nt ins (116) | 1nt del (138) | OK | OK | 1nt del (87) | 2nt del (53) | Stop (39); 1nt ins (213) | OK |
| *Phocoena sinus* (**Phocoenidae**) | Exon Not Found | 1nt ins (116) | 1nt del (138) | OK | OK | 1nt del (87) | 2nt del (53) | Stop (39); 1nt ins (213) | OK |
| *Monodon Monoceros* (**Monodontidae**) | Exon Not Found | 1nt ins (116) | 1nt del (142) | OK | OK | 1nt del (87) | 2nt del (53) | Stop (39); 1nt ins (213) | OK |
| *Delphinapterus leucas* (**Monodontidae**) | Exon Not Found | Stop (51); 1nt ins (116) | 1nt del (121); 1nt del (138); 1nt del (182) | OK | OK | 1nt del (87) | 2nt del (53) | 1nt ins (213) | OK |
| *Lipotes vexillifer* (Lipotidae) | Exon Not Found | Stop (51); 1nt ins (116) | 1nt del (121); 1nt del (138); 1nt del (182) | OK | OK | 1nt del (87) | 2nt del (53) | 1nt ins (213) | OK |
| *Physeter microcephalus* (**Physeteridae**) | Unknown (Incomplete Genomic Sequence) | Unknown (Incomplete Genomic Sequence) | Unknown (Incomplete Genomic Sequence) | Unknown (Incomplete Genomic Sequence) | Unknown (Incomplete Genomic Sequence) | 1nt del (87) | OK | Stop (246) | OK |
| *Balaenoptera acutorostrata scammoni* (Balaenopteridae) | Exon Not Found | OK | 2nt del (10) | OK | OK | 1nt del (87); 1nt del (134) | OK | Stop (246) | OK |

**Legend:**

ins – insertion

del – deletion

stop – in-frame premature stop codon

(xxx) – (coordinate of the annotated deleterious mutation relative to the reference exon)
